# CDK4/6 Inhibition Uncovers Subtype-Specific Vulnerabilities and Immune-Related Responses in Esophageal Squamous Cell Carcinoma

**DOI:** 10.1101/2025.09.12.673056

**Authors:** Fabiana Moresi, Diego Japon Ruiz, Matteo Serra, Marta Avalos Moreno, Eloine Garcia, Katia Coulonval, Andrea Pavesi, Benjamin Beck, Xavier Bisteau

## Abstract

Esophageal squamous cell carcinoma (eSCC) is a highly aggressive malignancy with poor prognosis and limited therapeutic options. Although immune checkpoint inhibitors such as nivolumab, have shown clinical benefit, particularly in patients with high PD-L1 expression, this subgroup represents only a small fraction of eSCC cases. CDK4/6 inhibitors such as palbociclib have only been tested as second-line agents in eSCC, often in combination with EGFR inhibitors, with minimal benefit.

Our study evaluates palbociclib as a first-line therapy in treatment-naive eSCC models. Using a panel of 22 eSCC cell lines with integrated multi-omics and phenotypic assays, we identified three response subtypes, resistant, delayed and arrested, correlated to Rb-pathway status. Interestingly, in delayed responders, palbociclib induced replication stress, DNA damage, and unprotected micronuclei enriched for cGAS, triggering activation of interferon-stimulated genes. Consistent with this, palbociclib enhanced immune cell infiltration in delayed eSCC spheroids within a preclinical vascularized 3D microfluidic system.

Our study demonstrates that first-line palbociclib treatment unmasks intrinsic vulnerabilities in the CDK4/6-Rb axis and triggers innate immune activation in molecularly defined eSCC. Using a translationally relevant 3D vascularized microfluidic system, we provide evidence that early CDK4/6 inhibition not only stall cancer cell growth but also promotes immune cells recruitment.

In conclusion, our study identifies palbociclib as a viable first-line therapeutic candidate in selected eSCC patients and uncover its immunomodulatory potential.

**Significance:** These findings support further research into CDK4/6 inhibition as first-line treatment for eSCC and highlight its potential to influence the tumor immune microenvironment in ways that could improve responses to combination therapies.

## Introduction

Esophageal squamous cell carcinoma (eSCC), the predominant histological subtype of esophageal cancer worldwide (1), is associated with a 5-year survival rate below 20% and ranks among the top ten causes of cancer-related mortality globally (2).

Its highly heterogeneous mutational landscape contributes to poor therapeutic outcomes and complicates the adoption of novel alternative treatments (3). The current standard of care, surgical resection combined with neoadjuvant genotoxic therapy, provides only modest long-term benefit, with many patients ultimately experiencing disease relapse or progression. For advanced or metastatic eSCC, platinum-based chemotherapy remains the mainstay of treatment as the first-line approach (1).

More recently, immune checkpoint inhibitors such as nivolumab and pembrolizumab have been approved in combination with chemotherapy for selected patients, particularly those with elevated PD-L1 expression (4). However, this group represents a minority, and most eSCC tumors remain immunologically "cold" (5), limiting clinical benefit and underscoring the urgent need for alternative strategies.

Among the most frequently altered oncogenic pathways in eSCC, the cyclin D–CDK4/6– RB1 axis is frequently dysregulated through amplification of *CCND1*, homozygous deletion or epigenetic silencing of *CDKN2A*, and, less commonly, loss or mutation of *RB1* (6–8). These alterations collectively drive cell proliferation by accelerating G1/S phase transition, consequently creating a potential therapeutic vulnerability in eSCC. Three CDK4/6 inhibitors, like palbociclib, have been reported to provide a significant clinical benefit in the treatment of HR+/HER-breast cancer (9). This led to their approval as standard first-line therapies in that setting (10). However, in eSCC, CDK4/6 inhibitors have so far been evaluated exclusively in second-line approaches, like in combination with EGFR inhibitors under the ongoing clinical trial *NCT05865132*, and typically following prior cytotoxic therapies. In this context, their efficacy has been limited, likely due to acquired resistance mechanisms and a reduced dependency on CDK4/6 signaling (11).

Recently, CDK4/6 inhibition has been increasingly studied in earlier therapeutic windows for eSCC. Notably, the clinical trial *NCT06654297* is assessing the combination of palbociclib with the anti–PD-1 antibody camrelizumab in a neoadjuvant setting for resectable eSCC but results are still pending. To date, the immunomodulatory effects of palbociclib in eSCC remain largely unexplored, particularly as first-line option and within complex tumor-immune microenvironments.

Emerging evidence from other tumor types has reported that CDK4/6 inhibitors have unexpected additional effects beyond cell cycle arrest. These include modulation of DNA damage responses (12), induction of senescence (13), and stimulation of innate immune signaling (14). Notably, such effects can reshape the tumor microenvironment (TME), promoting a shift toward immune stimulation. For instance, in hepatocellular carcinoma, palbociclib has been shown to induce senescence-associated inflammatory signaling that enhances CD8⁺ T cell–mediated cytotoxicity (15). These findings suggest that early intervention with CDK4/6 inhibitors, prior to the establishment of resistance to other therapies, could not only uncover therapeutic vulnerabilities but also reprogram the TME to favor anti-tumor immune responses.

Despite mounting interest for CDK4/6 inhibition, no study has yet systematically profiled treatment-naive eSCC models to dissect both molecular and immunologic mechanisms across multiple cell lines. Prior work has focused primarily on single-cell line analyses, such as the radiosensitizing and cytostatic effects of palbociclib (16,17). While these studies highlight important therapeutic effects, none have combined multi-lines, multi-omics profiling in treatment-naive eSCC, which is the core contribution of our study.

Here, we performed a comprehensive multi-omics and phenotypic analysis of the response of 22 eSCC cell lines to palbociclib to identify molecular vulnerabilities and mechanisms that influence immune signals. This approach revealed three distinct response groups, (i) resistant, (ii) delayed, and (iii) arrested, correlating with Rb pathway status and cell cycle dynamics. Morphological and transcriptomic profiling of these cells unveiled context-specific micronuclei formation and elevated expression of chemokines and cytokines. Based on this classification, arrested models appear to respond well to palbociclib monotherapy, while delayed responders exhibited replication-associated DNA damage, accumulation of γH2AX foci, and partial activation of cytosolic DNA sensing pathways. This pattern is interesting, as immunogenic stress suggests potential benefit from immunotherapy combinations.

A preclinical 3D co-culture model, recapitulating key features of the tumor microenvironment, including perfusion, stromal interactions, and spatial organization (18), easily assessed and unveiled immune responses to palbociclib, indicating its utility as a platform to identify response patterns without the need for predefined biomarkers. In conclusion, palbociclib appears to be a viable first-line therapeutic candidate in molecularly defined eSCC and may be leveraged for its immunomodulatory potential.

## Materials and Methods

### Cell culture

A panel of 22 cell lines was used in this study (see Supplemental Table 1 for full information from Cellosaurus and culture parameters). The KYSE and TE cells series were respectively obtained from DSMZ (Brunswick, Germany) and from the cell bank of the RIKEN BRC (Tsukuba, Japan). These authenticated cell lines were passaged for fewer than 4 months. All cells were cultured in RPMI (Gibco RPMI medium 1640 1X; ref: 21875-034) or F12 medium (Gibco F12 nut mix 1X; ref: 21765-029) supplemented with 1% penicillin/streptomycin (Nacalai Tesque 09367-34) and fetal bovine serum (Gibco 26140-079) at concentrations ranging from 2% to 10%, according to the specific medium requirements for each cell line. Cells were maintained in a 5% CO_2_ atmosphere and a constant humidity.

### Clonogenic assay

Cells were seeded in 6-well plates at densities ranging from 2 × 10² to 4 × 10³ cells/well, depending on their respective doubling times. After 48 h, cultures were treated with either vehicle control (DMSO) or palbociclib (1 µM final concentration kinetic experiments or a gradient of concentrations 16nM-4000nM for IC50 experiments) for up to 18 days. The culture medium was refreshed every 3 days. At defined time points, cell monolayers were fixed in 10% neutral buffered formalin (Sigma-Aldrich) for 5 min at room temperature, rinsed with PBS, and stained with 1 mL of 0.05% crystal violet (Sigma-Aldrich) for 30 min under continuous slow agitation. Plates were washed four times by submersion in tap water and air-dried before image acquisition with a Sony Alpha 7 mirrorless camera. Colony confluency was quantified using the *ColonyArea* plugin in ImageJ (19). Obtained raw growth curves were normalized by scaling all values to the maximum confluency observed under DMSO treatment, yielding normalized area values (0–100%). Proliferation inhibition was quantified using the area under the curve (AUC) from both conditions, computed via a trapezoidal rule in R (version 4.3.3) using the package “pracma” (version 2.4.4). A normalized “response score” was calculated as

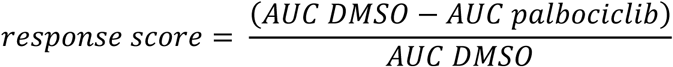

reflecting the proportional reduction in growth due to treatment, independent of each line’s inherent growth capacity. The three phenotypic patterns were defined based on a corrected response score, calculated by subtracting the standard deviation (SD) of the response score from the mean response score for each cell line. Cell lines were classified as: Resistant if the corrected score was ≤ 0; Delayed if the corrected score was > 0 and < 0.90; and Responsive if the corrected score was ≥ 0.90. This correction accounted for experimental variability, ensuring robust classification independent of replicate noise. For IC₅₀ determination, normalized dose–response curves were fitted to a four-parameter logistic regression model using the drc package in R (v 3.0.1) (20), from which IC₅₀ values were derived for each cell line.

### Immunoblotting

Cells were lysed by manual scraping in Laemmli 1x sample buffer supplemented with protease and phosphatase inhibitors [60µg/mL Pefabloc, 4 mM Naf, 1µg/mL Leupeptine, 100µM Vanadate] before denaturation at 95°C for 5 minute, snap frozen in liquid nitrogen and stored at −80 °C until further use. Protein content was evaluated by paper method (21). Equal amounts (10-30 μg) of whole cell extract proteins were separated by SDS-PAGE (ranging 6-12%) and transferred onto PVDF membrane (Immobilon®-P IPHV00010, 0.45 µm). Membranes were blocked in blocking buffer (5% non-fat dry milk {Bio-Rad, 1706404} in 0.1% Tween20 TBS) at room temperature before incubation with primary antibodies (see Supplemental Table 2 for full references) diluted in blocking buffer overnight at 4°C. Membranes were incubated with appropriate HRP-conjugated secondary antibodies (diluted in blocking buffer) at room temperature before successive washes and their revelation using enhanced chemiluminescence (WesternLightning® Plus ECL and Western Sirius [Advansta K-12043-D10]).

For 2D-gel electrophoresis, cells were lysed using a chaotropic buffer (7M urea, 2M thiourea). Protein extracts were subjected to isoelectric focusing (IEF) using immobilized pH gradient (IPG) strips (pH 5–8 for CDK4 {ReadyStrip™ IPG Strips #1632018, BioRad}). Following IEF, protein separation was completed by SDS-PAGE as described in previously established protocols (22,23). Chemiluminescence detections were captured with a Vilber-Lourmat Solo7S camera. The electrophoretic pattern of CDK4 generated by this approach has been previously characterized (23,24).The most basic isoform (spot 1) corresponds to the unmodified CDK4, while the most acidic (spot 3) represents CDK4 phosphorylated at threonine 172, confirmed using phospho-specific antibodies.

## RNA-seq Data Processing and Analysis

### RNA sequencing

Total RNA was isolated from cell lines treated with 1 µM Palbociclib or DMSO for 6 days using E.Z.N.A.® MicroElute® Total RNA Kit according to the manufacturer’s protocol. RNA yield and purity were assessed using a Fragment Analyzer 5200 (Agilent Technologies, Massy, France). One hundred nanograms of total RNA was used to construct indexed cDNA libraries using the NEBNext Ultra II Directional RNA Library Prep Kit for Illumina (New England Biolabs, Ipswich, MA, USA), according to the manufacturer’s instructions. The multiplexed libraries were sequenced on an Illumina NovaSeq 6000 system using a 200 Cycles S2 flow cell. Paired-end reads were aligned to the human reference genome GRCh38 using STAR, and transcript annotation was based on Homo_sapiens.GRCh38.90.gtf from Ensembl (ftp.Ensembl.org). Gene-level counts were generated using HTSeq and normalized as counts per 20 million reads (CP20M).

### Quality Control and Normalization

Raw RNA-seq counts were normalized using DESeq2’s median-of-ratios method to correct for library size and composition biases. Size factors were estimated using the estimateSizeFactors() function, and normalized counts were extracted with the counts(dds, normalized=TRUE) function (25). For downstream visualization and correlation analysis, a pseudo-count of 4 was added to avoid undefined log values, and data were transformed using log2(normalized count + 4). A pseudo-count of 4 was chosen empirically to stabilize variance in low-expression genes while minimizing the influence of technical noise. This log-transformed matrix was used to assess sample reproducibility via Pearson correlation.

### Gene Filtering and Differential Expression Analysis

Genes with low expression were filtered prior to analysis, retaining those with sufficient counts in at least two RNA-seq samples (see Supplemental Table 3 for data). Differential expression analysis was performed separately for each cell line using the DESeq2 package (v1.40.2). For each comparison (Palbociclib vs. DMSO), a DESeqDataSet object was created using a design formula ∼ treatment + replicate. After model fitting with DESeq, log2 fold changes were estimated and subsequently shrunk using the apeglm package (v1.24.0) implemented in the lfcShrink function to improve effect size estimation and reduce noise for lowly expressed genes (26). Genes were considered differentially expressed if they had an adjusted p-value (FDR) < 0.05, calculated using the Benjamini– Hochberg procedure, and an absolute log_2_ fold change > 1 (see Supplemental Table 3 for data). Results were visualized using heatmaps and volcano plots. Heatmaps displayed log2 fold changes of differentially expressed genes and were generated using the pheatmap or gplots packages. Volcano plots were created with ggplot2 and selected genes of interest were annotated using ggrepel to avoid label overlap.

### Gene Set Enrichment Analysis (GSEA)

Gene set enrichment analysis (GSEA) was performed using the fgsea R package (v. 1.34.2) (Korotkevich 2021). Genes were ranked by log₂ fold change from DESeq2 differential expression results. Enrichment was tested against curated MSigDB gene set collections, including Hallmark, C2 (curated gene sets: KEGG, Reactome, BioCarta), C5 (ontology gene sets: GO BP, CC, MF, and HPO) and C6 (oncogenic signatures). The fgseaMultilevel() function was used with parameters minSize = 10 and maxSize = 1000. Pathways with an adjusted p-value < 0.05 and a normalized enrichment score (NES) ≥ 1 or ≤ –1 were considered significantly enriched (see Supplemental Table 3 for data).

GSEA results were visualized with dot plots generated using ggplot2, summarizing NES and statistical significance across cell lines. To explore phenotype-specific enrichment trends, selected pathways of interest were further analyzed. A custom score (NES × – log10(p-value)) was computed to integrate enrichment magnitude and confidence. Selected KEGG pathways were visualized using the pathview package, with log₂ fold changes from DESeq2 as input. Statistical significance of pathway-level shifts was assessed using the gage package.

## BrdU Incorporation for Flow Cytometry

Cells were labelled with 5-bromo-2’-deoxyuridine (BrdU, sigma Aldrich B-5002) and 5-Fluoro-2′-deoxyuridine (FldU, sigma-adrich F-0503) by adding a mix of BrdU-FldU at a final concentration of 100 µM and 2 µM, respectively. Cells were incubated for 1 h or 24 h, depending on experimental conditions. After pulsed incorporation, adherent and floating cells were collected via trypsinization and centrifugation, washed with cold PBS, and fixed with ice-cold methanol added under gentle vortexing and incubated for ≥30 min at 4 °C. Fixed cells were resuspended in PBS containing 1% BSA (sigma-aldrich A9647) and treated with 2 N HCl/0.5% Triton X-100 for DNA denaturation. Samples were then neutralized with 0.1 M sodium tetraborate buffer (pH 8.5) and resuspended in PBS-BSA 1%. For BrdU detection, cells were stained with a primary anti-BrdU antibody (see Supplemental Table 2 for full references) at room temperature, washed, and incubated with a secondary antibody (see Supplemental Table 2 for full references) in the dark. Following antibody staining, cells were washed and counter-stained with propidium iodide (PI, 50 µg/mL final, sigma-aldrich #537059) and RNase A (sigma-aldrich R6513, 20 µg/mL final) in PBS-BSA. Stained cells were filtered and analyzed on a BD LSR-Fortessa or X20 to assess DNA content and BrdU incorporation. Analysis of scatter plots were performed using FlowJo V10 and statistical analyses were performed using R (v4.3.3). Comparisons between treatment groups were conducted using a paired *t*-test (Welch correction applied, i.e., var.equal = FALSE) grouped by cell line. P-values were adjusted for multiple comparisons using the Benjamini-Hochberg (BH) method, and adjusted significance levels were annotated accordingly, significance was defined as *p* < 0.05.

### Immunofluorescence 2D

Cells were seeded on µ-Slide 8 Well Glass Bottom plates (Ibidi 80827-90) and cultured for at least 24 hours before treatment. After 2 washes with PBS 1X, cells were fixed with 10% formalin (Sigma HT501128-4L) or cold methanol for 10 minutes, depending on the downstream application (e.g., BrdU staining, lamin). For permeabilization, cells were treated with 0.1% Triton X-100 in PBS for 10 minutes at room temperature and washed intensively with PBS. Blocking was carried out with 1:20 diluted goat (Jackson ImmunoResearch 005-000-121), or horse (Invitrogen, 26050070) serum in PBS containing 0.3% BSA (PBS/BSA 0.3%) under slow agitation for 1 hour. After washes, diluted primary antibodies in PBS/BSA 0.3% (see Supplemental Table 2 for full information and references) were added and incubated overnight at 4 °C. After washes, diluted secondary antibodies in PBS/BSA 0.3% (see Supplemental Table 2 for full information and references) were incubated for 2 hours under slow agitation at RT, before washes with PBS/BSA 0.3%. Nuclear counter-staining was conducted with propidium iodide (PI, 25 µg/mL final, sigma-aldrich 537059) and RNase A (sigma-aldrich R6513, 20 µg/mL final) in PBS-BSA for >15 min at RT in the dark. Cells were washed with distilled water before imaging. Images were acquired using an Aurox LFC spinning disk confocal microscope. Depending on the scope of the experiment, three different objectives were used: 10×/0.3 dry (WD 5.5 mm), 20×/0.8 dry (WD 0.55 mm), and 63×/1.4 oil (WD 0.19 mm). Acquired CZI images were processed using QuPath (v.0.5.1) and ImageJ (v.1.51) software.

### Nuclei segmentation and γH2AX foci analysis

Image analysis was performed using a combination of ImageJ and QuPath software. Z-stack fluorescence images were first processed in ImageJ (v1.51) to generate a maximum intensity projection and imported into QuPath (v0.4.3) for further analysis. Nuclear segmentation was conducted using the StarDist 2D plugin (27) (version 0.4.0) with a custom Python model. After segmentation, nuclei were manually reviewed, and smaller objects (∼1/8 the average nuclear size) were annotated as micronuclei. For γH2AX foci quantification, a single fluorescence intensity threshold was determined in QuPath for each cell line across all conditions. Statistical analyses were performed using R (v4.3.3). To assess differences in foci distributions between treatment conditions (e.g., DMSO vs.

D6) across cell lines, a two-sample Kolmogorov–Smirnov (KS) test was conducted. For each cell line, distributions of foci count per nucleus were compared between conditions. In addition to standard KS statistics, a directional KS score was calculated to quantify both the magnitude and direction of change. This score was defined as:

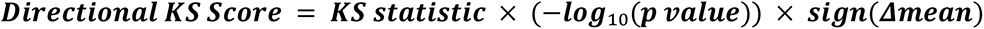

where *Δmean* is the difference in mean foci count between Day 6 and DMSO conditions. A positive score indicates an increase in foci upon day 6 treatment, while a negative score indicates a decrease. This composite metric captures statistical significance as well as the biological directionality and effect size, allowing for a more interpretable comparison across cell lines. To further evaluate condition-dependent differences in foci accumulation across cell lines, a generalized linear mixed model (GLMM) was fitted using a negative binomial distribution. Each model was run separately per cell line using the glmmTMB package (v1.1.11) (28) . For micronuclei frequency analysis, the percentage of micronucleated cells was computed per cell line and condition. As proportions were bounded between 0 and 1, a beta regression model was applied. All models were computed independently per cell line to account for inter-cell line variability in baseline damage (DMSO) and treatment response. Statistical significance was defined as *p* < 0.05.

### eSCC 3D model

#### Device seeding

HUVEC/TERT2 (EVERCYTE, CHT-006-0008) and LF/TERT309 (EVERCYTE, CHT-067-0309) fibroblasts were cultured in EGM-2 (Lonza, CC-3124) and DMEM/F12 (PAN Biotech, P04-41150) supplemented with 10% fetal bovine serum and 200 µg/mL G418 (InvivoGen, ant-gn5), respectively. Cells were trypsinized, counted, and resuspended at a final ratio of 6:1 (HUVEC:LF) in resuspension medium (RM) consisting of DMEM/F12 complete medium supplemented with Recombinant Human VEGF 165 Protein (VEGF-A, R&D, 293-VE), Normocure (InvivoGen, ant-noc-1), thrombin (Sigma-Aldrich, T7513-50UN), and collagen I (Corning, 354236). Tumor spheroids were generated in hanging drops and cultured for 4 days prior to use. Spheroids were harvested and transferred in a minimal volume of medium onto the lid of a 96-well plate. To prepare the gel mixture, 25 µL of the RM containing HUVECs, LFs, and spheroids was collected and mixed 1:1 with 25 µL of fibrinogen solution (3 mg/mL final concentration). The resulting 50 µL mixture was immediately dispensed into the central gel region of an OrganiX microfluidic chip (AIM Biotech, OGNXP-3EA), ensuring perpendicular pipetting to minimize bubble formation. Polymerization was allowed to proceed for 20 minutes at room temperature. Following polymerization, both media channels were hydrated with 40 µL of medium. An additional 230 µL and 330 µL of culture medium were added to the bottom and top left quadrants of the chip, respectively, to initiate directional flow across the gel. Devices were maintained at 37 °C in a humidified incubator under 5% CO₂. Culture media were changed every two days, and flow orientation was inverted daily for the first 4–5 days to promote vessel formation within the matrix. To assess vessel integrity and perfusability, Dextran-Texas Red dye (Thermo Fisher, D1830) was administered in one side channel of the chip between days 6 and 10, depending on the HUVEC batch. Dextran distribution and retention were monitored by live imaging using a Zeiss LSM 880 inverted confocal microscope.

#### PBMCs isolation

Peripheral blood samples were kindly provided by Giulia Adriani laboratory at A*STAR. Ethical approval for the collection of blood from healthy volunteers was granted by the institutional review board (Project No.: 201306-04. PBMCs were subsequently isolated according to the procedure described by Giustarini et al. (29)

### Immunofluorescence

OrganiX microfluidic devices containing vascularized tumor models were used, where tumor spheroids had been pre-labeled with CellTracker™ Green CMFDA Dye for live-cell tracking (Invitrogen, C7025). OrganiX microfluidic devices containing vascularized tumor models were washed three times with 1X PBS and fixed with 10% formalin overnight at 4 °C. Samples were subsequently rinsed with 1X PBS to remove excess fixative. For permeabilization, devices were incubated overnight at 4 °C with 0.1% Triton X-100 in PBS. To block nonspecific binding, devices were incubated overnight at 4 °C in a blocking buffer composed of 3% BSA (sigma-aldrich A9647) and 10% goat serum (Jackson ImmunoResearch 005-000-121) diluted in 0.1% Triton X-100 in PBS. Samples were then incubated in the dark with primary antibodies (see Supplemental Table 2) diluted in 0.5% BSA for 48h at 4 °C. Following primary antibody incubation, samples were washed in 0.1% BSA and incubated at 4°C overnight with secondary antibody (see Supplemental Table 2). After staining, devices were washed 1X PBS, followed by additional washes with MilliQ water to prepare the channels for optical clearing. Devices were imaged using a Zeiss LSM 880 inverted confocal microscope.

### Infiltration analysis via immunofluorescence

Multichannel fluorescence images were processed using FIJI/ImageJ (v1.53) and QuPath (v0.4.4). Z-stack images were converted into 2D projections using maximum intensity projection in FIJI and exported as TIFF files. In QuPath, tumor spheroids were manually segmented using the Luminicell channel and annotated as “Tumor”. CD31⁺ vessels were identified via pixel classification based on intensity thresholding, and CD45⁺ PBMCs were detected using the positive cell detection module with area and shape constraints. Spatial analysis was performed in R (v4.3.1). Distances between each CD45⁺ cell and the center of the spheroid were calculated and used to assign each cell to one of three radial compartments: spheroid (based on the segmented tumor area), proximity (a concentric region extending the spheroid radius by 40%), and periphery (extending the radius by 80%). PBMC density was defined as the sum of the areas of all immune cells within a given region, normalized to the total area of that compartment (µm²). Because spheroids were not always located precisely at the geometric center of the microfluidic device, this approach prevented bias in quantification, simply considering all segmented PBMCs without accounting for spatial distribution lead to overestimation of infiltration due to passive proximity rather than active recruitment.

### Statistical analysis

As density values were not normally distributed across conditions and compartments, we used the aligned rank transform (ART) ANOVA from the ARTool package in R (v. 0.11.2) to perform non-parametric factorial analysis (30) .

### Genomic Characterization and Integrative Clustering of eSCC Cell Lines

Genomic alteration data for esophageal squamous cell carcinoma (eSCC) cell lines were retrieved from the Cell Model Passport database and processed using R (v4.3.1). Copy number alteration (CNA) data were categorized based on the classification formula provided by Cell Model Passport:

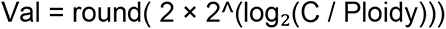

Where C represents the total copy number, Ploidy is the genome ploidy of the cell line, and Val is the discretized copy number value. CNAs were classified as follows:

- Val = 0 → *Deletion*
- Val = 1 → *Loss*
- Val = 2 → *Neutral*
- Val = 3 → *Gain*
- Val ≥ 4 → *Amplification*

Missing or “null” entries were imputed as Neutral. Data were reshaped into a numeric matrix with values ranging from 0 to 4 for each gene per cell line. Somatic mutations were binarized (1 = mutated, 0 = wild type. To explore molecular heterogeneity among the cell lines, we performed integrative clustering using the MOVICS (31) package (version 0.99.17). Normalized basal RNA-seq data (generated in-house), along with processed CNA and mutation data, were used as input. The optimal number of clusters was determined through Cluster Prediction Index and Gaps-statistics metrics using getClustNum() function. Clustering was then performed using the iClusterBayes method (32). Comprehensive heatmaps were generated using getMoHeatmap() with sample annotations including *RB* and *TP53* alteration status. Gene set variation analysis (GSVA) were performed with the function runGSVA() using curated MSigDB gene set Hallmark.

## Data Availability Statement

The sequence data generated in this study will be made publicly available in Gene Expression Omnibus (GEO) at reviewing (currently uploaded), and within the article and its supplementary data files. The CNA and mutation data were obtained from the Broad Institute’s Cancer Dependency Map project and were obtained from https://depmap.org/portal/data_page/?tab=customDownloads.

## Results

### Integrative profiling of eSCC cell lines uncovers distinct molecular subtypes and frequent alterations affecting cell cycle–associated genes

To evaluate whether esophageal squamous cell carcinoma (eSCC) is vulnerable to CDK4/6 inhibition as first line treatment, we first determined the molecular landscape of a panel of 22 eSCC cell lines that are frequently used as eSCC experimental models. Integrative multi-omics analysis, combining our generated data for bulk RNA sequencing with publicly available mutational profiling, and copy number alterations (CNA), captured the transcriptional and genomic heterogeneities of this cell panel. The unsupervised multi-omics clustering defined four distinct molecular subgroups (referred to as CS1–CS4), as depicted by the comprehensive heatmap in **Fig. 1A**, which reflects the global transcriptional and mutational profiles across the cell line panel. Differential gene expression assessment comparing each cluster of cell lines to the rest, further revealed distinct expression profiles for each cluster as displayed for top-upregulated genes (**Fig. 1B**). Gense GSVA based on Hallmark pathways highlighted distinct transcriptional programs across the four defined eSCC clusters (**Supplementary Fig. S1A**). CS1 appeared to display an upregulation for genes linked to immune and inflammatory pathways, including interferon-α and interferon-γ responses. In contrast, various cells in CS2 exhibited an upregulated mesenchymal and hypoxia-driven program, with marked enrichment of TGF-β signaling and epithelial–mesenchymal transition (EMT). CS3 and CS4 showed mixed phenotypes. CS4 separated from CS2 and CS3 clusters because of an upregulation of proliferative and cytokine-driven gene programs, with upregulation of MYC Targets, supporting cell cycle progression and biosynthesis, despite a downregulation of interferon response programs in comparison to CS1. Taken together, this multi-omics framework integrating expression, mutation, and CNA data enabled the identification of biologically distinct eSCC subgroups.

**Figure 1.**
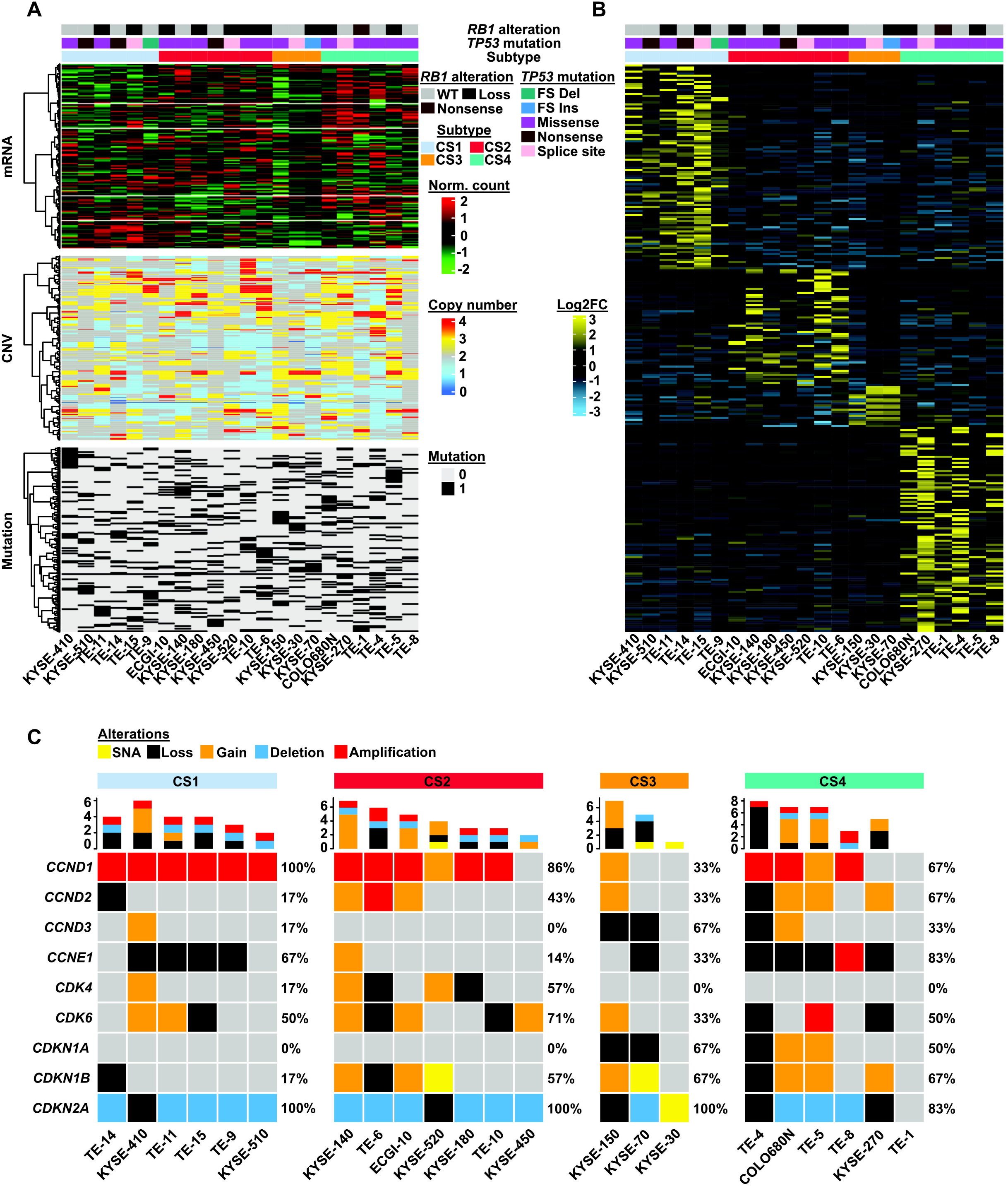
Mutational background of eSCC cell lines **(A)** Comprehensive multi-omics heatmap displaying RNA expression, copy number variation (CNA), and mutation data across 22 eSCC cell lines, used as input for integrative clustering using the MOVICS framework. Cell lines were classified into four molecular subtypes (CS1–CS4), as indicated*. TP53* and *RB1* mutation statuses are annotated above the heatmap. **(B)** Differentially expressed genes (DEGs) derived from the RNA-seq data are shown in a clustered heatmap across the CS1–CS4 subtypes, highlighting subtype-specific transcriptional signatures. **(C)** Cluster-specific OncoPrint visualization of alterations in genes related to the cell cycle machinery, including amplifications, deletions, gains, losses, and point mutations (SNA). The top barplot indicates the number of alterations per cell line, while the right percentages reflect the frequency of gene alterations within each cluster.

We further wondered whether this panel of eSCC cell lines was recapitulating genomic alterations identified in eSCC tumors and their frequencies. The integrated genomic characterization of esophageal carcinoma reported by the Cancer Genome Atlas Research network’s study (8) described key biological pathways that are frequently affected by mutation or CNAs, including, among others, the cell cycle, cell differentiation, or chromatin remodeling. Oncogenic alterations found in the panel of eSCC cells aligned with numerous hallmark mutations observed in primary tumors. The profiling of frequently altered genes involved in squamous cell differentiation (*TP63*, *SOX2*, *NOTCH1*) and chromatin remodeling (*SMARCA4*, *ARID1A*, *KMT2D*, *KDM6A*) (**Supplementary Fig. S1B**) revealed frequent alterations. Whereas chromatin remodelers were more commonly lost or mutated with a variable frequency among clusters, lineage regulators *SOX2* and *TP63* displayed frequent copy number gains in eSCC cell lines. Beside these variable alterations, all 22 cell lines harbored *TP53* mutations, primarily missense (**Fig. 1A–B**) like primary tumors. Immunoblotting analysis further indicated a complete loss of p53 protein expression for several cell lines and revealed changes in its size in most cases, potentially affecting its function (**Supplementary Fig. S2A**). This convergence suggests that dysfunctional or absent p53 is a unifying feature across the panel.

The cell cycle pathway appeared the most affected, particularly within the cyclin D– CDK4/6–Rb axis. Amplification of *CCND1*, deletion, loss of heterozygosity and mutation of *CDKN2A*, and copy number gains of *CDK4* and *CDK6* were prevalent across the 4 clusters (**Fig. 1C**). Although the loss of one *RB1 allele* was identified across several cell lines (**Fig. 1A**), two cell lines among the CS4 lacked the expression of the Rb protein (**Supplementary Fig. S2A**). The TE-1 cell line carries a nonsense mutation in *RB1*, whereas, for KYSE-270, despite presenting no Rb expression, no genomic alterations for *RB1* were found. The latter may reflect epigenetic silencing or an undetected structural alteration affecting *RB1*.

Loss of Rb expression was accompanied with high expression of p16 and an increased expression of cyclin E for KYSE-270. Expression of CDK2 barely varied between the entire panel, inversely to the abundance of CDK4 and CDK6 in several cell lines (**Supplementary Fig. S2A**). Expression of cyclin D1 displayed a basal expression in all eSCC cell lines but KYSE450, while some higher expression in various cell lines of the TE series. The phosphorylation of CDK4 on threonine 172 (pT172) is key for D-type cyclin/CDK4 complex activity, pushing cell cycle entry through the phosphorylation of Rb (33,34). Levels of phosphorylation on T172 was reported in various studies to correlate with the response to CDK4/6 inhibitors and has been proposed as a functional biomarker of CDK4/6 pathway dependency (24,35,36). Using bidimensional electrophoresis and immunoblotting of CDK4 (**Supplementary Fig. S2B**), we detected marked signals for pT172 CDK4 in most Rb-proficient cell lines such as KYSE-180, KYSE-410, or TE-10. No signal for pT172 CDK4 was observed in Rb-deficient lines (KYSE-270 and TE-1), similar to previous observations in other Rb-deficient cancers (24,35,36). However, several cell lines like KYSE-150, KYSE-450, TE-6, or TE-4 presented a minimal signal of phosphorylated CDK4.

Taken together, these observations mirror key mutational features reported in the TCGA eSCC cohort (8,37,38), further supporting the relevance of this panel of cell lines as a faithful model of patient-derived tumor biology. Oncogenic alterations, particularly in cell cycle-associated genes together with the elevated phosphorylation status of CDK4 in most Rb-proficient cells, reinforce the idea that eSCC cells would be sensitive to CDK4/6 inhibitors like palbociclib. However, the highlighted molecular heterogeneity among eSCC cell lines caution us that responses to CDK4/6 inhibitors may be unlikely uniform across all cell models. Moreover, it remains uncertain whether molecular subgroup classification alone is sufficient to reliably predict therapeutic response.

### Diverse response to palbociclib in eSCC cell lines reveal three main patterns

We therefore assessed the response of the cell line panel to CDK4/6 inhibition using palbociclib, as the most characterized CDK4/6 inhibitor (**Fig. 2A**). Across the entire panel, palbociclib sensitivity varied broadly, with IC₅₀ values ranging from sub-micromolar to greater than 4 µM, reflecting a spectrum from highly sensitive to intrinsically resistant cell lines (**Fig. 2B, C**). Consistent with the known requirement for functional Rb in mediating CDK4/6 inhibitor efficacy, Rb-deficient lines such as KYSE-270 and TE-1 exhibited no changes in their proliferation upon palbociclib treatment. Their BrdU incorporation remained high, indicating continued S-phase progression despite drug exposure (**Fig. 2D, E**) and no difference was observed between proliferation curves over 18 days of continuous drug treatment (**Fig. 2F, G**).

**Figure 2.**
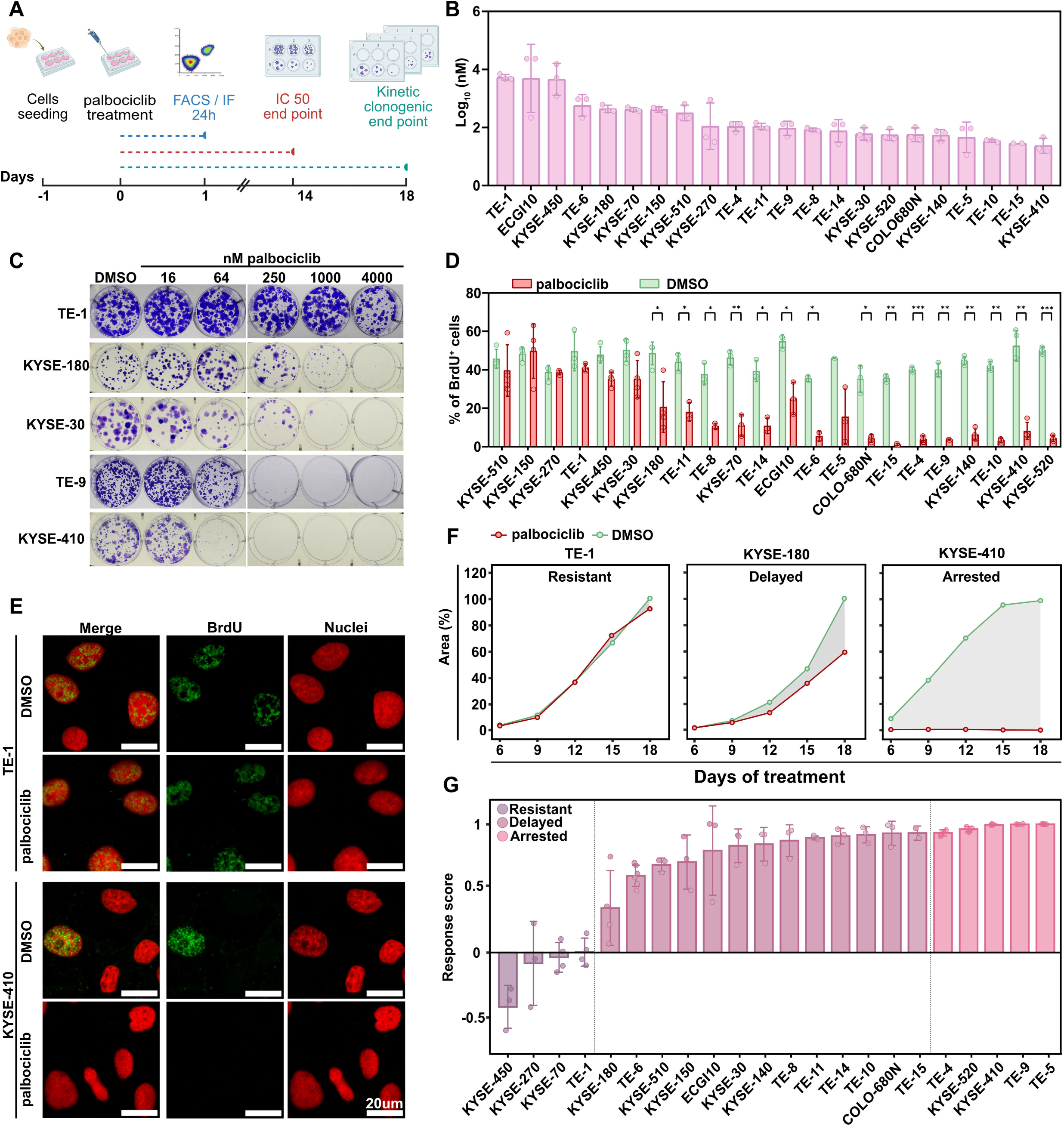
ESCC cells heterogeneously respond to CDK4/6 inhibition over short and long term. **(A)** Schematic overview of the experimental workflow, including short-term BrdU incorporation assays, long-term colony formation, and proliferation kinetics under continuous palbociclib treatment. **(B)** IC50 values for palbociclib (µM) calculated from colony formation assays after 14 days of treatment across 22 eSCC cell lines. Dots represent IC50 from each biological replicate (n=3); error bars display ±SD. **(C)** Representative images from colony formation assays in selected cell lines treated with increasing concentrations of palbociclib (16–4000 nM) for 14 days. **(D)** Quantification of BrdU incorporation (% BrdU⁺ cells) after 24 hours of treatment with 1 µM palbociclib or DMSO, measured by flow cytometry across the panel of eSCC lines. Data are shown as combined bar/dot plots, with individual replicate values, mean, and ±SD. significance was determined using a paired Student’s *t*-test followed by Benjamini-Hochberg (BH) multiple testing correction. **(E)** Representative immunofluorescence images of TE-1 and KYSE-410 cells showing BrdU (green) and nuclear (red) staining after 24-hour treatment with DMSO or 1 µM palbociclib; scale bar=20 µm. **(F)** Representative proliferation curves for selected eSCC cell lines showing differential responses to palbociclib: resistant (TE-1), delayed (KYSE-180), and arrested (KYSE-410) phenotypes, monitored every 3 days by crystal violet staining over 18 days. Grey areas indicate the difference of the area under each curve (AUC) between conditions. **(G)** Response scores representing relative changes in proliferation under palbociclib vs. DMSO, calculated from area under the curve (AUC) over an 18-day period as (AUC_DMSO – AUC_Palbociclib)/AUC_DMSO; data are shown as combined bar/dot plots, with individual replicate values (≥3), mean, and ±SD. Significance levels: p < 0.05 (*), p < 0.01 (**), p < 0.001 (***).

In contrast, most Rb-proficient cell lines like KYSE-410 or TE-9 showed a pronounced sensitivity to palbociclib, with clear cytostatic responses. Such cell lines displayed a flattened proliferation curve over prolonged treatment, leading to a substantial drop in cumulative area under the curve (AUC) compared to the control condition, and consequently presenting a high response score (**Fig. 2F, G**). BrdU incorporation in these sensitive models dropped significantly (**Fig. 2D, E**). Interestingly, an intermediate response pattern was observed in other Rb-proficient cell lines like KYSE-180 and TE-6. Under palbociclib treatment, both cell lines displayed a slower proliferation (**Fig. 2F, G)**. This pattern was characterized as a delayed proliferation, below a 90% threshold of proliferation reduction over 18 days (this pattern hereafter is referred to “Delayed; see the methodology section for complete criteria and calculation). Concomitant to this delay, BrdU incorporation was only partially reduced for both cell lines, parallel to a higher IC50 than most sensitive cells (**Fig. 2D**), suggesting the notion that more intricate mechanisms such as activated upstream pathways or combined oncogenic alterations generate an overall shift towards a more lenient restriction checkpoint, consequently modulating the sensitivity in this subgroup.

These data highlight once more that palbociclib efficacy is primarily associated with *RB1* status but temper the notion that all Rb-proficient cells respond similarly to CDK4/6 inhibition although our results predict a global sensitivity of eSCC to CDK4/6 inhibition with a minimal number of intrinsic resistance. These observations, however, also unveil a less comprehensive trend where palbociclib only slow down the proliferation of various cell lines. This trend may modulate additional observed phenotypes like DNA damage responses, induction of senescence or stimulation of innate immune signaling while the dynamics of the cell cycle upon prolonged palbociclib exposure would provide novel avenues to decipher the difference between proliferation patterns.

### The dynamic Rb pathway modulation defines palbociclib response in eSCC cell lines

To understand the differences of behavior between “delayed” and “arrested” subgroups, we examined the dynamics of cell cycle-related protein abundance by performing a time-course analysis over prolonged palbociclib exposure (**Fig. 3 & Supplementary Fig .3**). In KYSE-410, one of the most sensitive lines, levels of phosphorylated Rb rapidly dropped after one day, remaining low over the entire course despite a marginal rebound at Day 3. This behavior also appeared in other “arrested” lines like TE-4 and TE-5 but also in the delayed line TE-6. In contrast KYSE-180 (“delayed”) showed a partial reduction of Rb phosphorylation from Day 1 to Day 6, but the signal remained appreciable throughout. Notably, cyclin E did not decrease over the time course, and cyclin A/B changes were modest. This trend was also presented by other “delayed” lines like ECGI-10. As expected, because of their lack of Rb expression, TE-1 and KYSE-270 showed no detectable Rb signal at any time point, confirming their intrinsic resistance. Higher expression level of cyclin E in both these cell lines could explain their continuous proliferation, therefore relying on CDK2 activity. The increasing expression of cyclin E also helps to understand the higher proliferation of unexpected “resistant” cell lines KYSE-450 and KYSE-70, shifting their dependency toward CDK2 activity at a quick pace. In contrast, in arrested KYSE-410 and other lines such as TE-4 and TE-9, cyclin E levels remained low throughout the time course, with no appreciable increase, consistent with a stable G1 arrest and sustained suppression of E2F activity. For most “arrested” lines, cyclin A and cyclin B levels also rapidly declined, further supporting the maintenance of a G1 blockade.

**Figure 3.**
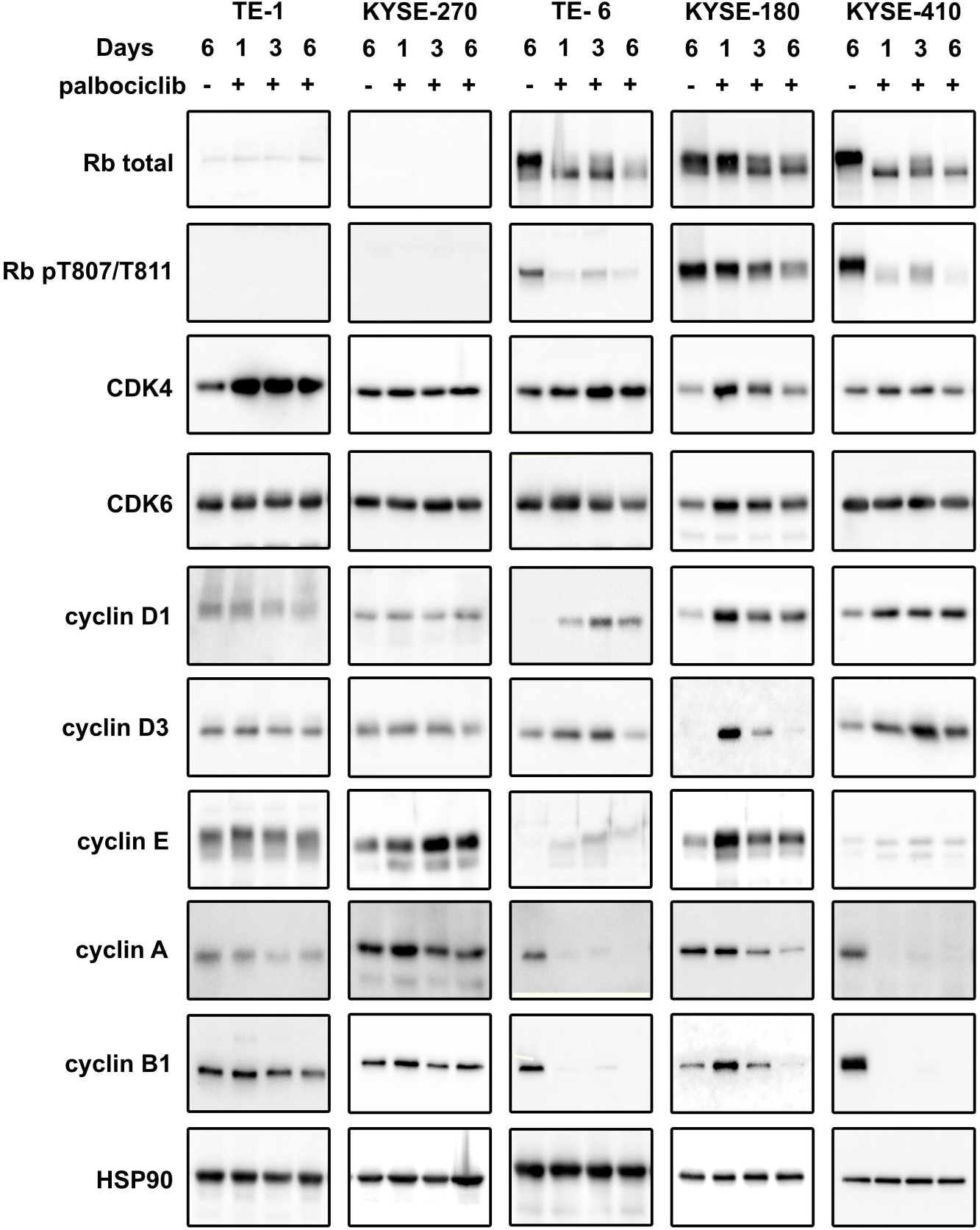
Protein expression dynamics following 6 days of palbociclib treatment. Western blot analysis on whole cell lysates collected after 6 days of continuous palbociclib treatment in representative eSCC cell lines. The panel includes phosphorylated Retinoblastoma protein (RB pS807/811), total Rb, CDK4, CDK6, cyclin D1, cyclin D3, cyclin A, cyclin E, and cyclin B1. HSP90 was used as a loading control.

These observations further support the importance of evaluating the functional status of Rb but these also indicate the complex regulations of kinase activities regulating the cell cycle engagement around the inhibition of Rb via its phosphorylation. The activating phosphorylation of CDK4 has appeared to be so far the best proxy to predict the responsiveness of cancer cells to CDK4/6 inhibitors like palbociclib. However, the dynamics of response observed for eSCC cell lines indicate an intricate situation where levels of regulators, particularly of cyclin E, allow cells to quickly adjust their proliferation. The consequences of these dynamics render difficult to comprehend the different phenotypes observed in multiple models upon CDK4/6 inhibition and need to be evaluated considering this heterogeneity and dynamics.

### Palbociclib induces time-dependent accumulation of unprotected micronuclei and activation of cytosolic DNA sensing in selected eSCC cell lines

Recent work by Crozier *et al*.(39) reported that p53-deficient cell lines accumulate nuclear abnormalities such as micronuclei following CDK4/6 inhibition. Since all 22 eSCC cell lines studied here exhibit *TP53* loss-of-function mutations or complete p53 loss (**Fig. 1A**), we investigated how the dynamics and the heterogeneous proliferation sensitivity to palbociclib were affecting the appearance of nuclear abnormalities.

Immunofluorescence images after 6 days of palbociclib-treatment revealed the appearance of micronuclei as illustrated for KYSE-180, inversely to palbociclib-resistant TE-1 cell line (**Fig. 4A**). Quantification of cells with micronuclei for all 22 lines (**Fig. 4B**) further established the statistically significant appearance and accumulation of such aberrant micronuclear bodies in 19 out of 22 cell lines. Both unresponsive cell lines to palbociclib TE-1 and KYSE-270 presented no significant difference between the basal control condition and 6 days of palbociclib treatment. Temporal analysis of micronuclei appearance in KYSE-180 (**Fig. 4C**) showed a steady accumulation over time, reaching the highest levels at Day 6 of palbociclib treatment. In contrast, TE-1 maintained high basal levels of micronuclei without increase in cell proportion over time, reinforcing its classification as intrinsically resistant. Importantly, the extent of micronuclei appearance correlated significantly with the proliferation response score upon palbociclib exposure across the cell line panel (**Fig. 4D**), with more sensitive cell lines displaying greater differences in percentage of cells with micronuclei. Inversely, resistant ones such as TE-1, KYSE-270, KYSE-450 and KYSE-70 clustered in the lower-left portion of the plot, indicating that micronuclei accumulation is tightly linked to palbociclib sensitivity.

**Figure 4.**
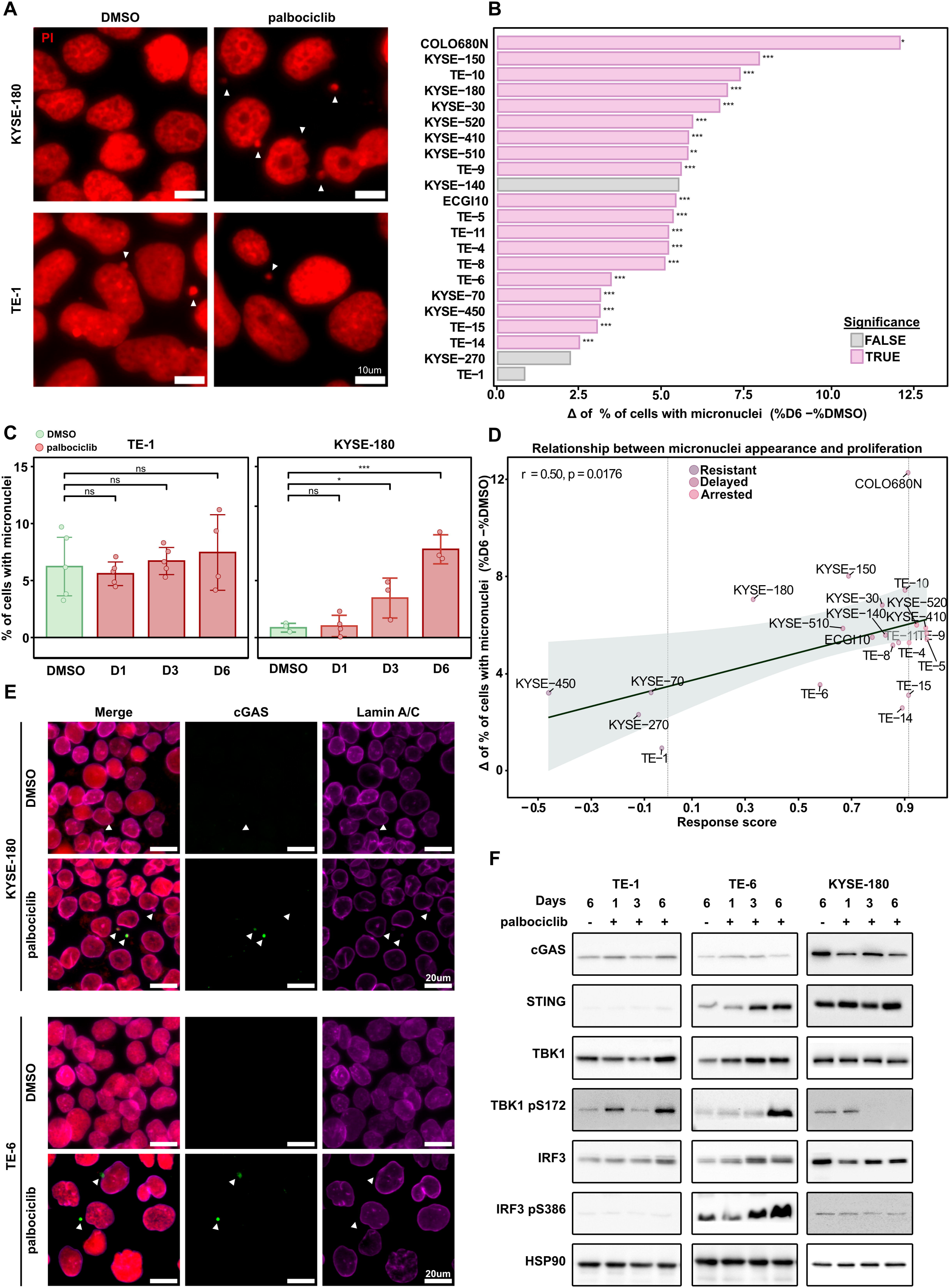
Palbociclib treatment induces micronuclei accumulation and activation of the cGAS–STING pathway in eSCC cell lines. **(A)** Representative immunofluorescence images showing micronuclei (white arrowheads) in two eSCC cell lines (TE-1 and KYSE-180) after 6 days of 1 µM palbociclib treatment or DMSO control. Nuclei were counterstained with propidium iodide (PI). Scale bar = 20 µm. **(B)** Quantification of micronuclei accumulation across 22 eSCC cell lines at day 6. Values represent the percentage difference in micronucleated cells (Δ micronuclei (%)) between DMSO and palbociclib (1 µM, Day 6) conditions. **(C)** Quantification of micronuclei appearance over time in TE-1 and KYSE-180 across 6 days of palbociclib treatment (1 µM). Each dot represents an independent biological replicate; bars show means ± SD. **(D)** Correlation analysis between palbociclib growth inhibition (response score) and Δ micronuclei (%). Each point represents an individual cell line. **(E)** Representative immunofluorescence images of micronuclei (PI) co-stained with Lamin A/C (violet) and cGAS (green) in TE-6 and KYSE-180 after 6 days of palbociclib treatment. Scale bars = 20 µm. **(F)** Western blot analysis of cGAS–STING pathway components in three eSCC cell lines (TE-1, TE-6, KYSE-180) treated for 1, 3, or 6 days with 1 µM palbociclib or DMSO. HSP90 was used as a loading control. Significance levels: p < 0.05 (*), p < 0.01 (**), p < 0.001 (***).

Because different studies have reported that micronuclei can induce immunogenic signals upon rupture of their nuclear envelope, thereby exposing their DNA content to cytosolic sensors such as cGAS (40), we investigated whether the micronuclei observed upon prolonged exposure in eSCC cells would trigger cytosolic DNA sensors. We assessed nuclear integrity and cytosolic DNA sensing through co-immunostaining of Lamin A/C and cGAS, respectively (**Fig. 4E**). Representative images from KYSE-180 and TE-6, two cell lines presenting significant micronuclei appearance upon palbociclib treatment, revealed that while nuclear envelope rupture (indicated by Lamin A/C negative staining) is required for cGAS recruitment, not all ruptured micronuclei necessarily recruit cGAS, additional chromatin states may influence this outcome. Quantification of the different categories of micronuclei over time (**Supplementary Fig. S4A**) further supported this observation, presenting a statistically significant increase of delaminated DNA (Lamin-negative, cGAS-positive) in the cytoplasm of both cell lines following prolonged palbociclib treatment.

To evaluate whether the signaling downstream of cGAS is activated, we looked at the cGAS–STING–TBK1–IRF3 axis by immunoblotting across all 22 eSCC cell lines (**Supplementary Fig. S4B**), with representative examples highlighted in the main figure (**Fig. 4F**). TE-6, which exhibited elevated proportion of cGAS-positive micronuclei (**Fig. 4E**), also showed a progressive rise in phosphorylated TBK1 (pS172) and IRF3 (pS386) over the 6-day treatment, indicating full activation of the pathway. In contrast, KYSE-180, despite showing prominent cGAS-positive micronuclei, failed to activate the canonical STING–TBK1–IRF3 axis, raising the possibility that cGAS may be channelled toward alternative signaling outputs. In addition to TE-6, other cell lines such as TE-4 and TE-14 also showed activation of STING pathway effectors, as evidenced by increased levels of p-TBK1 and p-IRF3 (**Supplementary Fig. S4B**). These data confirm that canonical cGAS–STING signaling is activated in a subset of eSCC models following palbociclib treatment, though not universally, and its consequence needs to be assessed.

### Palbociclib-Induced Micronucleation Triggers Immune-Related Transcriptional Reprogramming

Since we observed that palbociclib treatment induced the accumulation of unprotected, micronuclei, we next investigated the transcriptional consequences of this phenomenon. RNA sequencing was performed on all 22 eSCC cell lines after 6 days of treatment, corresponding to the time point when the micronuclei burden peaked. Gene Set Enrichment Analysis (GSEA) revealed consistent downregulation of hallmark cell cycle pathways (e.g., E2F targets, G2M checkpoint, Mitotic spindle), particularly in palbociclib-sensitive lines, while immune-related signatures, including interferon alpha/gamma response, IL6, JAK–STAT3 signaling, and TNFα via NF-κB, were enriched in a subset of cell lines (**Fig. 5A**).

**Figure 5.**
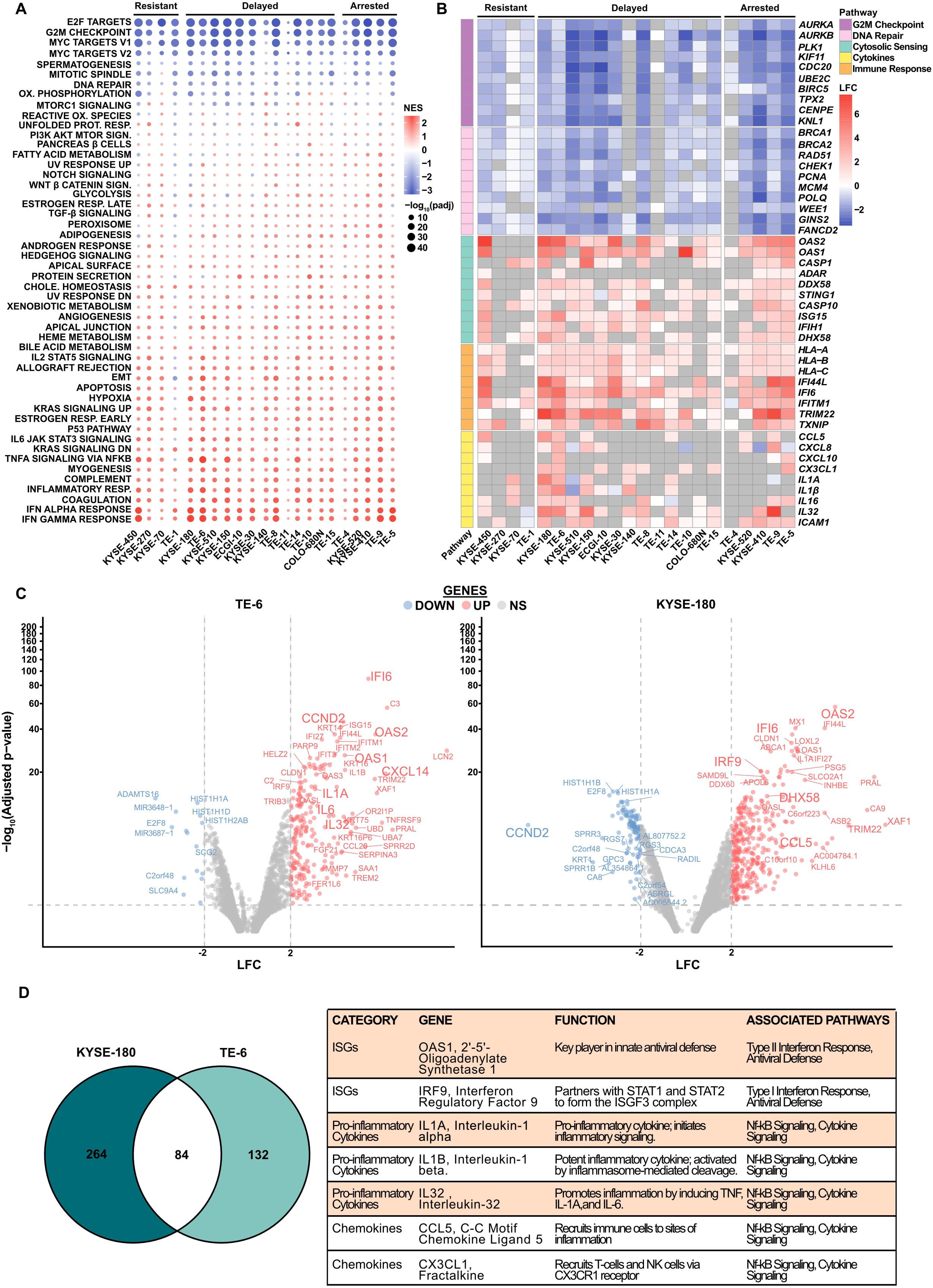
Transcriptional response to CDK4/6 inhibition reveals phenotype-specific pathway rewiring. **(A)** Gene Set Enrichment Analysis (GSEA) performed on RNA-seq data from 22 esophageal squamous cell carcinoma (eSCC) cell lines after 6 days of palbociclib treatment. The dot plot shows the Normalized Enrichment Score (NES) of Hallmark pathways for each cell line. Dot color indicates NES direction (red = positive enrichment, blue = negative enrichment), and dot size reflects statistical significance (p-value). **(B)** Heatmap of selected gene sets involved in cell cycle, immune response, and cytosolic DNA sensing pathways, stratified by cell line and phenotype. **(C)** Volcano plots showing gene-level differential expression (DEG) between untreated and palbociclib-treated TE6 (left) and KYSE-180 (right) cell lines. Genes were classified as upregulated if log_2_ fold change ≥ 2 and FDR < 0.05, or downregulated if log_2_ fold change ≤ –2 and FDR < 0.05. Non-significant genes were labeled as "NS". Red and blue dots represent significantly upregulated or downregulated genes, respectively. **(D)** Overlap of significantly upregulated genes between KYSE-180 and TE-6 visualized by Venn diagram (left). Table (right) highlights a selection of shared palbociclib-induced genes involved in inflammatory or innate immune pathways.

To explore these trends further, we focused on curated gene sets related to cell cycle arrest, immune signaling, DNA repair, and cytosolic DNA sensing **(Fig. 5B**). This visualization highlights robust upregulation of interferon-stimulated and inflammatory genes, such as *OAS1*, *OAS2*, *ISG15*, *IFIH1*, *DDX58*, *CASP1*, *CASP10*, and *CXCL10*, many of which are known downstream targets of cytosolic DNA sensing and type I interferon signaling (41). These transcriptional signatures were particularly evident in models such as TE-6 and KYSE-180, both of which showed cGAS-positive micronuclei formation. Notably, resistant lines such as TE-1 and KYSE-270 did not exhibit similar transcriptional changes, consistent with their lack of micronuclei induction and cGAS-STING pathway activation. Volcano plots of TE-6 and KYSE-180 revealed strong upregulation of shared immune response genes including *OAS1*, *OAS2*, *IFI6*, *TRIM22*, *CXCL14*, and *CCL5*, reinforcing the link between micronuclei formation and innate immune activation (**Fig. 5C**). Overlap analysis identified a common subset of significantly induced genes in both KYSE-180 and TE-6 (**Fig. 5D**), suggesting convergence on innate immune transcriptional programs downstream of cytosolic DNA sensing. These results indicate that palbociclib not only enforces cell cycle arrest in susceptible eSCC cells but also trigger inflammatory transcriptional responses, particularly in models predisposed to micronuclear rupture, offering new perspectives on its immunomodulatory potential.

### Palbociclib induces DNA damage in a subset of eSCC cell lines

To further investigate the upstream events contributing to micronuclei formation and innate immune activation, we assessed the induction of DNA damage in palbociclib-treated eSCC cell lines using γH2AX as a marker. Immunofluorescence staining revealed an increase in nuclear γH2AX foci in depicted lines after 6 days of treatment, with marked differences depending on the sensitivity to the drug (**Fig. 6A**). For instance, TE-1 cells, which were resistant to palbociclib, displayed abundant γH2AX foci under basal (DMSO) control conditions but did not exhibit any notable increase upon treatment. In contrast, KYSE-180, characterized by a delayed proliferation phenotype, showed a clear rise in γH2AX foci following palbociclib exposure, suggesting ongoing DNA damage accumulation during partial cell cycle progression.

**Figure 6.**
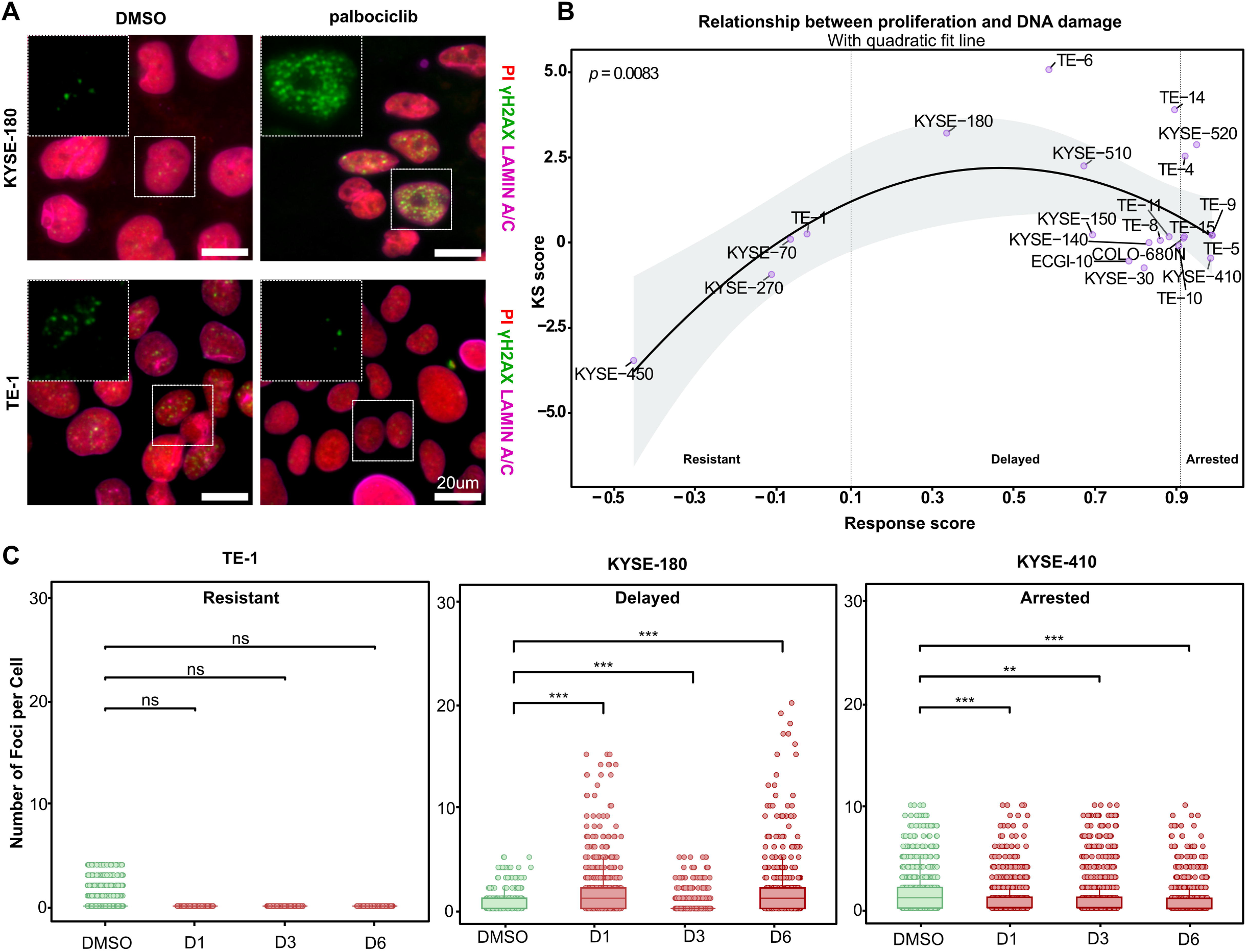
Palbociclib treatment induces DNA damage in selective cell lines eSCC cell lines. **(A)** Representative immunofluorescence images of two selected eSCC cell lines (TE-1 and KYSE-180) treated for 6 days with 1 µM palbociclib or DMSO. Cells were stained for γH2AX (green), nuclei (PI, red) and Lamin A/C (pink). White arrowheads indicate the appearance of micronuclei. Scale bar=20 µm. **(B)** Correlation between DNA damage (measured as Kolmogorov–Smirnov [KS] score from γH2AX foci distributions) and palbociclib growth response score (delta AUC, Figure 2). Each point represents an individual cell line; the quadratic fit line depicts the trend between DNA damage accumulation and treatment sensitivity. **(C)** γH2AX foci distribution in three representative eSCC lines (TE-1, KYSE-180, KYSE-410) treated with DMSO, 1 µM palbociclib for 1, 3, or 6 days. Individual data points represent the number of foci per cell; lines indicate medians. Statistical significance was assessed per cell line using a generalized linear mixed model (GLMM) with a negative binomial distribution. Significance levels: p < 0.05 (*), p < 0.01 (**), p < 0.001 (***).

A global correlation analysis across the 22 cell lines revealed an inverted U-shaped relationship between palbociclib response (as defined by Response score in **Fig. 2G**) and γH2AX-based DNA damage scores, referred as KS score (**Fig. 6B**). Specifically, cell lines with intermediate sensitivity, such as KYSE-180 and TE-6, exhibited the highest levels of γH2AX accumulation, suggesting persistent replication stress in the context of incomplete cell cycle arrest. In contrast, intrinsically resistant lines (e.g., TE-1) and strongly sensitive lines (e.g., KYSE-410) showed relatively low levels of DNA damage.

γH2AX foci quantification further supported this interpretation (**Fig. 6C**). KYSE-410, which displayed a strong inhibition of its proliferation, exhibited a progressive decrease in γH2AX foci over the 6-day treatment period. KYSE-180, in contrast, showed a gradual accumulation of DNA damage, consistent with its incomplete cell cycle arrest leading to replication stress. TE-1, as expected for a resistant line, maintained a steady γH2AX signal throughout and showed no significant induction upon treatment.

Collectively, these results suggest that palbociclib induces DNA damage in a subset of eSCC cell lines, particularly those that fail to fully arrest in G1.

### Palbociclib promotes immune cell recruitment in a 3D microfluidic model of eSCC

To determine whether the innate immune signaling observed in palbociclib-treated eSCC models translates into enhanced immune cell engagement, we adapted our findings to a 3D microfluidic setting using the OrganiX devices. eSCC spheroids (TE-1, TE-6, KYSE-180) were generated via hanging drops and embedded within a perfusable endothelial network (**Fig. 7A**). Before drug treatment, vascular integrity and perfusion were verified by injecting RFP-conjugated 70 kDa dextran. Notably, the presence of eSCC spheroids did not impair the formation or functionality of the endothelial network, as comparable perfusion was observed in chips with and without tumor spheroids (**Fig. 7B**). Once vessel functionality was confirmed, spheroids were treated with palbociclib for six days. Drug efficacy was first validated in 3D by measuring spheroid size. TE-1 remained resistant, showing continued growth despite treatment, whereas TE-6 and KYSE-180 spheroids displayed moderate inhibition, mirroring their “delayed” response phenotype previously observed in 2D monolayer cultures (**Fig. 7C**).

**Figure 7.**
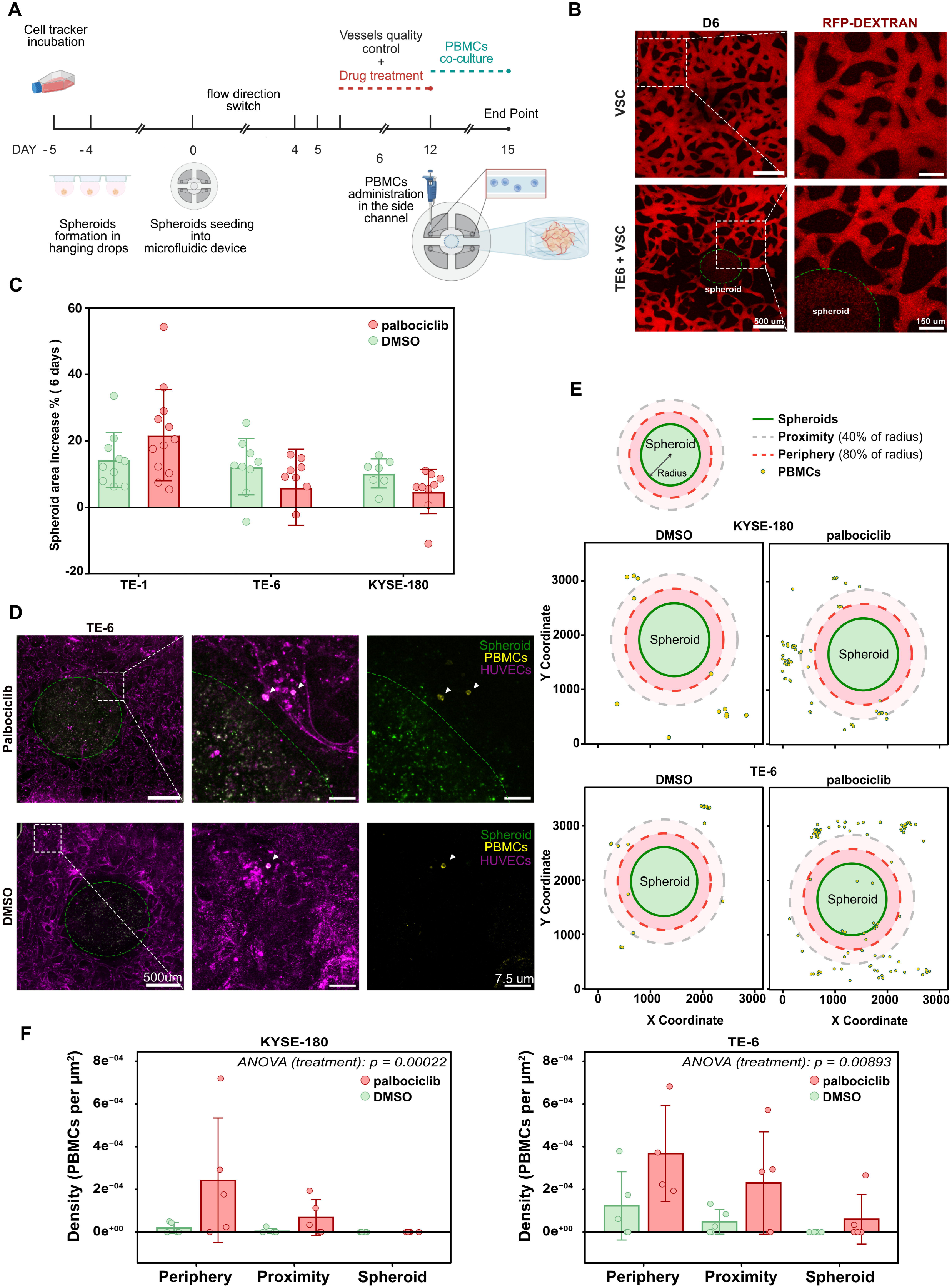
Palbociclib modulates immune cell infiltration in a 3D vascularized eSCC microfluidic model. **(A)** Schematic timeline of the experimental workflow. eSCC spheroids were formed in hanging drops and embedded into microfluidic chips. Vascular structures (VSC) were allowed to form before initiating treatment with 1 µM palbociclib from Day 0 to Day 6. **(B)** Representative images showing vascular perfusion using RFP-conjugated 70 kDa dextran, acquired 6 days after microfluidic seeding. Top: vessels alone; bottom: vessels co-cultured with TE-6 spheroids. Scale bars = 500 µm (overview), 150 µm (inset). **(C)** Quantification of spheroid area increase (Day 6 vs. Day 0), expressed in percentage. Each dot represents an individual microfluidic device; bars indicate mean ± SD across three eSCC cell lines. **(D)** Representative immunofluorescence images of TE-6 spheroids co-cultured with PBMCs following 6 days of treatment with Palbociclib or DMSO. HUVECs are shown in magenta, PBMCs in yellow, and spheroids in green. Insets highlight increased PBMC proximity to spheroids in Palbociclib-treated conditions compared to DMSO. Scale bars = 500 µm (overview), 7.5 µm (inset). **(E)** Computational mapping of PBMCs spatial distribution in the OrganiX device after PBMCs co-culture. PBMC positions (cyan dots) are plotted relative to the spheroid center for palbociclib (left) and DMSO (right) conditions. Coordinates were extracted from QuPath segmentations of the immunofluorescence images and processed in R. Red and gray dashed circles indicate the spheroid, proximity, and periphery regions used for density quantification. **(F)** Quantification of PBMC spatial density across three concentric radial regions: spheroid, proximity, and periphery. Density was calculated as the PBMC area normalized to the surface area of each region. Statistical significance was assessed using aligned rank transform (ART) ANOVA; p-values for treatment effects are indicated.

To evaluate immune cell recruitment, PBMCs were introduced into the side channel of the device on Day 12 and allowed to flow through the vasculature. After 3 days of co-culture, immunofluorescence imaging revealed enhanced proximity of PBMCs to palbociclib-treated spheroids compared to DMSO controls (**Fig. 7D**). We defined artificial concentric regions (spheroid, proximity, periphery) to normalize spatial variability. Computational mapping of PBMC positions relative to the tumor core showed clear enrichment in the spheroid and peri-spheroid regions under palbociclib (Proximity and Periphery, **Fig. 7E**). Quantitative analysis of PBMC density across three concentric radial zones confirmed a statistically significant increase in immune infiltration in both immunologically active cell lines TE-6 and KYSE-180 (**Fig. 7F**). These findings support the notion that palbociclib can reshape the tumor microenvironment to favor immune cell engagement in selected eSCC models, particularly those displaying features of innate immune activation at transcriptomic level.

## Discussion

While platinum-based chemotherapy remains the standard of care of first-line therapy against eSCC (42), palbociclib has only been studied in clinics in pretreated or refractory settings, with limited clinical benefit. Here, we present the first comprehensive evaluation of palbociclib as a potential first-line therapy in eSCC, using a panel of 22 cell lines characterized by diverse oncogenic alteration, the vast majority of which are derived from patients who had not received prior cytotoxic therapy. The only exceptions are KYSE-510 and KYSE-150, where patients had been previously treated with cisplatin and radiotherapy. The panel of cells used in this study exhibited substantial molecular heterogeneity, including the presence of distinct transcriptional and genomic subtypes, mirroring the diversity observed in primary eSCC tumors (8,38,43,44). The diversity among cell lines allowed us to uncover three distinct response types to palbociclib, (i) resistant, (ii) delayed, and (iii) arrested. These results highlight how relying on limited number of cells can obscure significant variations in drug sensitivity, reinforcing the need for subtype-informed, multi-model systems in preclinical drug development. Integrative clustering of the 22 eSCC cell lines allowed defining four molecularly distinct subgroups (CS1–CS4) with divergent transcriptional and genomic features. While this derived from *in vitro* models, these clusters parallel patient-derived eSCC subtypes recently described by Jiang *et al.* (46), who integrated transcriptomic, histological, and single-cell data to define four tumor-intrinsic categories (Differentiated, Immunogenic, Metabolic, and Stemness). The comparison between both classification systems highlighted striking overlaps, underscoring both the biological consistency of eSCC heterogeneity and the translational relevance of cell line models.

Our CS1 cluster displayed upregulation of inflammatory pathways, similarly to the Immunogenic subtype described by Jiang et al. Although cell lines inherently lack a native immune microenvironment, the transcriptional enrichment we observe suggests a retained immunomodulatory potential in these models. CS2 cells exhibited some enrichment of TGF-β signaling and epithelial–mesenchymal transition (EMT), consistent with a stem-like phenotype. Frequent *SOX2* copy number gains further reinforce this dedifferentiated state. This profile mirrors the Stemness subtype in patient tumors, which Jiang et al. associated with poor prognosis. CS3 was distinguished by selective activation of the PI3K–AKT–mTOR pathway. This phenotype resonates with elements of Jiang *et al.*’s Metabolic subtype, which was enriched in drug metabolism pathways. Both classifications converge on the notion of metabolic rewiring as a tumor survival strategy. Importantly, our CS3 cluster contains several resistant or delayed-response cell lines to palbociclib (e.g., KYSE-70 and KYSE-150), suggesting that it may serve as tractable model to combine with therapeutic approach linked to metabolic pathways or PI3K– mTOR inhibition. CS4 was characterized by MYC target activation, cell cycle progression, and IL6–JAK–STAT3/IL2–STAT5 signaling. This proliferative/cytokine-driven program does not map one-to-one onto a single Jiang’s subtype but shares features with both the Differentiated subtype and, to some extent, the Immunogenic subtype, given its cytokine engagement. Importantly, CS4 also included cell lines with Rb-deficient, a feature critical for predicting response to CDK4/6 inhibition.

Both frameworks underscore the non-uniformity of eSCC and the existence of reproducible transcriptional states across tumors and cell models. Key oncogenic alterations, including *CCND1* amplification, *CDKN2A* deletion, *SOX2/TP63* copy number changes and universal TP53 loss, were observed in our cell line panel and align with the mutational spectra in TCGA and Jiang *et al.*’s cohorts (8,46).

While these shared genomic alterations highlight the consistency of eSCC hetereogenity across models, their direct implications for CDK4/6 pathway vulnerability require deeper analysis. To this end, we investigated candidate biomarkers predictive of palbociclib sensitivity.

At the molecular level, among the evaluated biomarkers, *RB1* status emerged as the most definitive determinant of intrinsic resistance. For instance, TE-1 and KYSE-270, both Rb-deficient and lacking CDK4 phosphorylation on Thr172, showed no phenotypic response to palbociclib, reinforcing the concept that functional Rb is essential for CDK4/6 inhibitor efficacy (45). In this context, loss of *RB1* serves as a “master negative marker” for patient exclusion from palbociclib-based therapies, as its absence reliably predicts resistance regardless of other pathway components.

Conversely, sensitivity to palbociclib was generally associated with the co-occurrence of three features: intact *RB1*, amplification of *CCND1* and active CDK4 signaling, as marked by phosphorylation on Thr172. However, our findings also emphasize the limitations of relying on single biomarkers for predicting treatment outcomes. For instance, KYSE-450, despite being *RB1*-proficient, lacked both *CCND1* amplification and pT172 CDK4 and was resistant to palbociclib. Likewise, KYSE-180 and TE-6, both delayed responders, retained *RB1* and presented *CCND1* amplification while harboring CDK4 loss, potentially perturbing cyclin D/CDK stoichiometry and favoring compensatory activation of CDK6– cyclin D3 and CDK2–cyclin E pathways. These observations suggest that coordinated pathway activity, rather than the presence of isolated markers, dictates treatment response, a concept supported by similar findings in HR+/HER2– breast cancer (46).

Functionally, palbociclib induced the accumulation of micronuclei in most cell lines except the resistant group, with a peak around Day 6. Delayed responders such as TE-6 and KYSE-180 displayed increased γH2AX-positive micronuclei, indicating DNA damage. While we did not directly assess the source of these damages, previous studies have suggested that CDK4/6 inhibition can lead to replication stress and fork stalling (39,47), which may underline this phenotype. In contrast, arrested models accumulated micronuclei without elevated γH2AX levels, suggesting that these structures may arise through mechanistically distinct, non-replication-associated processes. Indeed, senescent or prolonged G1 arrested cells have been shown to accumulate chromatin aberrations and lamina stress, which may predispose to micronuclear formation via chromatin remodeling and nuclear fragmentation (48).

These observations were further supported by our quantitative analysis of γH2AX levels and their correlation with treatment response. Notably, palbociclib-induced DNA damage followed an inverted U-shaped trend across the cell line panel: resistant cells (e.g., TE-1) and arrested cells (e.g., KYSE-410, TE-4, TE-5) displayed minimal γH2AX induction and intermediate (delayed) responders (e.g., KYSE-180, TE-6) exhibited the highest levels of DNA damage accumulation. This pattern is consistent with two distinct behaviors: resistant cells, which maintain proliferation despite treatment and therefore do not accumulate DNA damage from CDK4/6 blockade, and arrested cells, which avoid damage by exiting the cell cycle and remaining in G1. In contrast, partially responsive models exhibited persistent DNA damage throughout treatment, a phenomenon similarly described by Crozier *et al.*, who demonstrated that prolonged CDK4/6 inhibition impairs origin licensing and increases DNA damage (40). In these contexts, subthreshold CDK4/6 inhibition may allow aberrant S-phase entry, replication fork stalling, and ultimately DNA damage, as reported in breast cancer (39) and bone marrow (49). In addition, recent work has demonstrated that CDK4/6 inhibitors can impair homologous recombination by promoting PARP1 degradation in Rb intact non-small cell lung cancer (50).

Significantly, although palbociclib treatment induced widespread micronuclei formation across most eSCC cell lines, the downstream immunologic signals varied substantially. For instance, COLO-680N exhibited the highest frequency of micronuclei by Day 6 post-treatment; however, these structures failed to recruit cGAS, indicating that micronuclei alone are insufficient to initiate innate immune sensing. While prior studies, such as those by Jiang *et al*. (12), have reported that palbociclib-induced micronuclei robustly activate the cGAS–STING pathway in melanoma and CRC models, our data highlight a more nuanced outcome in eSCC. These findings underscore the tumor-type and context-specific nature of CDK4/6 inhibitor–mediated immune remodeling. One possible explanation lies in the intrinsic differences in micronuclear composition and integrity, such as chromatin compaction, nuclear envelope integrity, or histone modifications, which can limit the accessibility of DNA to cytosolic sensors (51). One possibility is that micronuclei arising in cells with low proliferation—or those in arrested states—retain more compact chromatin and intact histone association, features known to diminish cGAS recruitment. Indeed, MacDonald *et al.* demonstrated that transcriptionally active micronuclei are less prone to cGAS binding, and Takaki *et al.* showed in their model that histone-bound DNA within micronuclei can impede cGAS activation. While these studies suggest potential barriers to cGAS engagement, our findings indicate that when micronuclei are formed under replicative conditions, they may remain accessible to cGAS, supporting the idea that the proliferative context of micronuclei formation influences their immunogenic potential (51,52).

Conversely, in cell lines such as KYSE-180 and TE-6, micronuclei were positive for cGAS, but only few of them like TE-6 proceeded to activate the canonical STING–TBK1–IRF3 signaling cascade. In KYSE-180, despite robust cGAS recruitment, TBK1 downstream effectors remains unphosphorylated. This suggests that cGAS localization is necessary but not sufficient to trigger the canonical pathway response, indicating that, in certain cellular contexts, cGAS may signal through noncanonical pathways, as reported in organ fibrosis (53). These could include NF-κB activation or engagement of a senescence-associated secretory phenotype (SASP), highlighting alternative routes by which palbociclib may modulate the tumor immune microenvironment (54).

Our findings highlight two distinct levels of immunologic heterogeneity across eSCC models: (i) differences in micronuclear competence for cGAS recruitment, and (ii) divergence in downstream immune signaling despite cGAS positivity. They underscore the need to consider not only the frequency but also the qualitative properties of micronuclei, such as chromatin state and nuclear envelope integrity, when evaluating the immunomodulatory potential of CDK4/6 inhibitors. Moreover, while cGAS–STING remains the best-characterized mediator of micronuclei-induced innate immunity (55), other sensors, such as AIM2, IFI16, or DDX41, may contribute to context-dependent immune responses (56,57). Further work will be needed to disentangle the relative contribution of these parallel sensing mechanisms in palbociclib-treated eSCC.

These findings suggest that palbociclib can elicit cytosolic DNA sensing responses in a subset of p53-deficient eSCC models, but the downstream immune signaling outcomes remain highly context dependent. The ability of certain lines to link micronuclei accumulation with cGAS–STING activation supports an immunogenic dimension of CDK4/6 inhibition and may provide a rationale for combinatorial strategies with immunotherapies, particularly in delayed responders with partial cell cycle escape and cytosolic DNA exposure.

The FDA’s recognition of organ-on-chip technologies under the 2.0 Modernization Act underscores their value as clinically actionable preclinical models (58). To validate our findings, we utilized OrganiX, a 3D vascularized microfluidic co-culture platform (18), and established the first vascularized esophageal squamous cell carcinoma model. Unlike traditional 2D models, our 3D model in the OrganiX platform recapitulates tumor architecture, perfusion, and immune cell access, enabling dynamic assessment of PBMC infiltration in response to treatment (59). Notably, we observed increased immune recruitment in palbociclib-responsive spheroids. This aligns with emerging data in other tumor types showing CDK4/6 inhibition can enhance tumor immunogenicity (60,61). Altogether, our findings suggest that early intervention with palbociclib may not only inhibit proliferation in a subset of eSCC models but also reprogram the tumor microenvironment toward a more immunostimulatory state. This opens new avenues for therapeutic synergy with immune checkpoint inhibitors, particularly in delayed responders who show partial cell cycle escape, replication stress, and cytosolic DNA sensing activation. Moreover, combining CDK4/6 inhibitors with agents targeting DNA damage response pathways, such as PARP or ATR inhibitors, may further amplify this phenotype as shown in recent studies across other tumor types (62).

The integration of multi-omics profiling, phenotypic characterization, and microfluidic co-culture provides a comprehensive framework for stratifying eSCC responses. Importantly, our classification of treatment-naïve cell lines into resistant, delayed, and arrested subtypes offers a translationally actionable strategy for guiding therapeutic decisions. Arrested models, marked by strong CDK4/6 dependency and minimal DNA damage, may benefit from palbociclib monotherapy. Conversely, delayed responders, despite initial suppression, exhibit features suggestive of immunogenic stress and may be prime candidates for rational immunotherapy combinations.

In this context, microfluidic-based 3D platforms like OrganiX could be further developed as robust preclinical tools for prospective functional testing and patient stratification. For instance, Li et al. recently established a lung cancer organoid, PBMC co-culture system (“GLI” model) that accurately reflected patient responses to anti–PD-1 immunotherapy in a manner predictive of clinical outcomes (63). Similarly, OrganiX could be used to test sensitivity of patient-derived eSCC spheroids to palbociclib, and even palbociclib plus immunotherapy, directly *ex vivo*. By observing PBMC infiltration and drug response in real time, clinicians could discern responders versus non-responders without relying solely on static genomic biomarkers.

Altogether, these findings lay the foundation for personalized treatment strategies in eSCC and may help reposition palbociclib from a second-line, post-chemotherapy setting to a first-line precision therapy in selected patient subgroups identified through functional testing rather than static biomarkers.

## Acknowledgements

We thank the FACS and LiMiF core facility for the help. We thank Anne Lefort and Frederick Libert for the sequencing performed at the Brussels Interuniversity Genomics High Throughput Core (www.brightcore.be). We thank Pierre Roger for support and discussions, Virginie Imbault for the lab management and discussions, and all present and past members of the Bisteau’s and Beck’s labs for fruitful discussions.

## Author Contributions

X.B. and F.M. conceptualized the study; F.M, D.J.R, E.G., K.C and X.B. performed experiments, analyzed, and interpreted data; F.M. & A.M. processed and analyzed publicly omics data; F.M. & M.S performed correlation analysis; F.M., B.B. and X.B wrote the paper; X.B. B.B. & A.P. supervised, provided resources and funding acquisition for the paper; and all authors revised and approved the manuscript.

---

**Supplementary Figure S1.** Mutational background of eSCC cell lines **(A)** Gene Set Variation Analysis (GSVA) of Hallmark gene sets across 22 eSCC cell lines. Heatmap displays pathway activity scores, highlighting heterogeneity in signaling programs relevant to proliferation, differentiation, metabolism, and immune response. Cell lines are grouped according to the MOVICS-derived clusters (CS1–CS4). **(B)** OncoPrint visualization showing genomic alterations in genes associated with squamous cell differentiation and chromatin remodeling complexes. The plot displays amplifications, deletions, gains, losses, and point mutations across the 22 eSCC cell lines, grouped by the MOVICS-derived clusters (CS1–CS4). Percentages indicate the proportion of altered cell lines per gene within each cluster.

**Supplementary Figure S2.** Mutational background of eSCC cell lines **(A)** Western blot analysis of key components of the cyclin D-CDK4/6–Rb pathway across 22 eSCC cell lines in proliferative untreated condition. Protein lysates from three breast cancer cell lines (MDA-MB-231, MDA-MB-468, and HCC1806) were included as reference controls. **(B)** 2D Western blotting of total CDK4 and its activating phosphorylation at threonine 172 (pT172) in representative untreated eSCC cell line.

**Supplementary Figure S3.** Protein expression dynamics following 6 days of Palbociclib treatment (extended panel). Western blot analysis on whole cell lysates collected after 6 days of continuous palbociclib treatment in 17 eSCC cell lines. The panel includes phosphorylated Retinoblastoma protein (Rb pS807/811), total Rb, CDK4, CDK6, cyclin D1, cyclin D3, cyclin A, cyclin E, and cyclin B1. HSP90 was used as a loading control. TE-1, KYSE270, TE-6, KYSE-180 and KYSE-410 cell lines are shown in the Figure 3 panel (A-B) as representative cell lines.

**Supplementary Figure S4.** Extended analysis of micronuclei composition and cGAS–STING pathway activation across eSCC cell lines. |**(A)** Quantification of micronuclei status over time in two eSCC cell lines (TE-6 and KYSE-180) treated with 1 µM palbociclib or DMSO. Micronuclei were classified as cGAS-positive (cGAS⁺), Lamin A/C-negative (Lamin⁻). Shown are the percentages of cGAS⁺ and Lamin⁻ micronuclei across days 1, 3, and 6. **(B)** Western blot analysis of cGAS–STING pathway activation across the full panel of 19 eSCC cell lines treated with 1 µM palbociclib or DMSO for 1, 3, or 6 days. TE-1, TE-6, and KYSE-180 cell lines are shown in Figure 4. HSP90 was used as a loading control.

## References

1. Obermannová R, Alsina M, Cervantes A, Leong T, Lordick F, Nilsson M, et al. Oesophageal cancer: ESMO Clinical Practice Guideline for diagnosis, treatment and follow-up ⋆. Ann Oncol. 2022;33:992–1004.

2. Matz M, Valkov M, Šekerija M, Luttman S, Caldarella A, Coleman MP, et al. Worldwide trends in esophageal cancer survival, by sub-site, morphology, and sex: an analysis of 696,974 adults diagnosed in 60 countries during 2000-2014 (CONCORD-3). Cancer Commun. 2023;43:963–80.

3. Liu Z, Zhao Y, Kong P, Liu Y, Huang J, Xu E, et al. Integrated multi-omics profiling yields a clinically relevant molecular classification for esophageal squamous cell carcinoma. Cancer Cell. 2023;41:181–195.e9.

4. Lu Y, Wang W, Wang F. Clinical benefits of PD-1 inhibitors in specific subgroups of patients with advanced esophageal squamous cell carcinoma: a systematic review and meta-analysis of phase 3 randomized clinical trials. Front Immunol. 2023;14:1171671.

5. Zheng Y, Chen Z, Han Y, Han L, Zou X, Zhou B, et al. Immune suppressive landscape in the human esophageal squamous cell carcinoma microenvironment. Nat Commun. 2020;11:6268.

6. Testa U, Castelli G, Pelosi E. The Molecular Characterization of Genetic Abnormalities in Esophageal Squamous Cell Carcinoma May Foster the Development of Targeted Therapies. Curr Oncol. 2023;30:610–40.

7. Song Y, Li L, Ou Y, Gao Z, Li E, Li X, et al. Identification of genomic alterations in oesophageal squamous cell cancer. Nature. 2014;509:91–5.

8. Kim J, Bowlby R, Mungall AJ, Robertson AG, Odze RD, Cherniack AD, et al. Integrated genomic characterization of oesophageal carcinoma. Nature. 2017;541:169–75.

9. Lin D-C, Wang M-R, Koeffler HP. Genomic and Epigenomic Aberrations in Esophageal Squamous Cell Carcinoma and Implications for Patients. Gastroenterology. 2018;154:374–89.

10. Martínez-Jañez N, Ezquerra MB, Sanchez LMM, Carrasco FH, Torres AA, Morales S, et al. First-line therapy with palbociclib in patients with advanced HR+/HER2− breast cancer: The real-life study PALBOSPAIN. Breast Cancer Res Treat. 2024;206:317–28.

11. Karasic TB, O’Hara MH, Teitelbaum UR, Damjanov N, Giantonio BJ, d’Entremont TS, et al. Phase II Trial of Palbociclib in Patients with Advanced Esophageal or Gastric Cancer. Oncol. 2020;25.

12. Fan H, Liu W, Zeng Y, Zhou Y, Gao M, Yang L, et al. DNA damage induced by CDK4 and CDK6 blockade triggers anti-tumor immune responses through cGAS-STING pathway. Commun Biol. 2023;6:1041.

13. Lee DH, Imran M, Choi JH, Park YJ, Kim YH, Min S, et al. CDK4/6 inhibitors induce breast cancer senescence with enhanced anti-tumor immunogenic properties compared with DNA-damaging agents. Mol Oncol. 2024;18:216–32.

14. Chaikovsky AC, Sage J. BEYOND THE CELL CYCLE: ENHANCING THE IMMUNE SURVEILLANCE OF TUMORS VIA CDK4/6 INHIBITION. Mol Cancer Res. 2018;16:molcanres.0201.2018.

15. Cai X, Yin G, Chen S, Tacke F, Guillot A, Liu H. CDK4/6 inhibition enhances T-cell immunotherapy on hepatocellular carcinoma cells by rejuvenating immunogenicity. Cancer Cell Int. 2024;24:215.

16. Qin W-J, Su Y-G, Ding X-L, Zhao R, Zhao Z-J, Wang Y-Y. CDK4/6 inhibitor enhances the radiosensitization of esophageal squamous cell carcinoma (ESCC) by activating autophagy signaling via the suppression of mTOR. Am J Transl Res. 2022;14:1616–27.

17. Wang J, Li Q, Yuan J, Wang J, Chen Z, Liu Z, et al. CDK4/6 inhibitor-SHR6390 exerts potent antitumor activity in esophageal squamous cell carcinoma by inhibiting phosphorylated Rb and inducing G1 cell cycle arrest. J Transl Med. 2017;15:127.

18. Adriani G, Pavesi A. The OrganiX microfluidic system to recreate the complex tumour microenvironment. Nat Rev Immunol. 2024;24:307–307.

19. Guzmán C, Bagga M, Kaur A, Westermarck J, Abankwa D. ColonyArea: An ImageJ Plugin to Automatically Quantify Colony Formation in Clonogenic Assays. Plos One. 2014;9:e92444.

20. Ritz C, Baty F, Streibig JC, Gerhard D. Dose-Response Analysis Using R. Plos One. 2015;10:e0146021.

21. Minamide LS, Bamburg JR. A filter paper dye-binding assay for quantitative determination of protein without interference from reducing agents or detergents. Anal Biochem. 1990;190:66–70.

22. Coulonval K, Bockstaele L, Paternot S, Dumont JE, Roger PP. The cyclin D3-CDK4-p27kip1 holoenzyme in thyroid epithelial cells: activation by TSH, inhibition by TGFβ, and phosphorylations of its subunits demonstrated by two-dimensional gel electrophoresis. Exp Cell Res. 2003;291:135–49.

23. Bockstaele L, Coulonval K, Kooken H, Paternot S, Roger PP. Regulation of CDK4. Cell Div. 2006;1:25.

24. Raspé E, Coulonval K, Pita JM, Paternot S, Rothé F, Twyffels L, et al. CDK4 phosphorylation status and a linked gene expression profile predict sensitivity to palbociclib. Embo Mol Med. 2017;9:1052–66.

25. Love MI, Huber W, Anders S. Moderated estimation of fold change and dispersion for RNA-seq data with DESeq2. Genome Biol. 2014;15:550.

26. Korotkevich G, Sukhov V, Budin N, Shpak B, Artyomov MN, Sergushichev A. Fast gene set enrichment analysis. bioRxiv. 2021;060012.

27. Schmidt U, Weigert M, Broaddus C, Myers G. Cell Detection with Star-convex Polygons. arXiv. 2018;265–73.

28. Brooks M E, Kristensen K, van Benthem K J, Magnusson A, Berg C W, Nielsen A, et al. glmmTMB Balances Speed and Flexibility Among Packages for Zero-inflated Generalized Linear Mixed Modeling. R J. 2017;9:378.

29. Giustarini G, Teng G, Pavesi A, Adriani G. Characterization of 3D heterocellular spheroids of pancreatic ductal adenocarcinoma for the study of cell interactions in the tumor immune microenvironment. Front Oncol. 2023;13:1156769.

30. Wobbrock JO, Findlater L, Gergle D, Higgins JJ. The aligned rank transform for nonparametric factorial analyses using only anova procedures. Proc SIGCHI Conf Hum Factors Comput Syst. 2011;143–6.

31. Lu X, Meng J, Zhou Y, Jiang L, Yan F. MOVICS: an R package for multi-omics integration and visualization in cancer subtyping. Bioinformatics. 2020;36:5539–41.

32. Mo Q, Shen R, Guo C, Vannucci M, Chan KS, Hilsenbeck SG. A fully Bayesian latent variable model for integrative clustering analysis of multi-type omics data. Biostatistics. 2018;19:71–86.

33. Bisteau X, Paternot S, Colleoni B, Ecker K, Coulonval K, Groote PD, et al. CDK4 T172 Phosphorylation Is Central in a CDK7-Dependent Bidirectional CDK4/CDK2 Interplay Mediated by p21 Phosphorylation at the Restriction Point. Plos Genet. 2013;9:e1003546.

34. Paternot S, Bockstaele L, Bisteau X, Kooken H, Coulonval K, Roger P. Rb inactivation in cell cycle and cancer: The puzzle of highly regulated activating phosphorylation of CDK4 versus constitutively active CDK-activating kinase. Cell Cycle. 2010;9:689–99.

35. Paternot S, Raspé E, Meiller C, Tarabichi M, Assié J, Libert F, et al. Preclinical evaluation of CDK4 phosphorylation predicts high sensitivity of pleural mesotheliomas to CDK4/6 inhibition. Mol Oncol. 2024;18:866–94.

36. Pita JM, Raspé E, Coulonval K, Decaussin-Petrucci M, Tarabichi M, Dom G, et al. CDK4 phosphorylation status and rational use for combining CDK4/6 and BRAF/MEK inhibition in advanced thyroid carcinomas. Front Endocrinol. 2023;14:1247542.

37. Zhao D, Guo Y, Wei H, Jia X, Zhi Y, He G, et al. Multi-omics characterization of esophageal squamous cell carcinoma identifies molecular subtypes and therapeutic targets. JCI Insight. 2024;9:e171916.

38. Liu M, An H, Zhang Y, Sun W, Cheng S, Wang R, et al. Molecular analysis of Chinese oesophageal squamous cell carcinoma identifies novel subtypes associated with distinct clinical outcomes. EBioMedicine. 2020;57:102831.

39. Crozier L, Foy R, Mouery BL, Whitaker RH, Corno A, Spanos C, et al. CDK4/6 inhibitors induce replication stress to cause long-term cell cycle withdrawal. Embo J. 2022;41:e108599.

40. Mackenzie KJ, Carroll P, Martin C-A, Murina O, Fluteau A, Simpson DJ, et al. cGAS surveillance of micronuclei links genome instability to innate immunity. Nature. 2017;548:461– 5.

41. Hornung V, Hartmann R, Ablasser A, Hopfner K-P. OAS proteins and cGAS: unifying concepts in sensing and responding to cytosolic nucleic acids. Nat Rev Immunol. 2014;14:521– 8.

42. Puhr HC, Prager GW, Ilhan-Mutlu A. How we treat esophageal squamous cell carcinoma. ESMO Open. 2023;8:100789.

43. Alexandrov LB, Kim J, Haradhvala NJ, Huang MN, Ng AWT, Wu Y, et al. The repertoire of mutational signatures in human cancer. Nature. 2020;578:94–101.

44. Liu W, Cui Y, Liu W, Liu Z, Xu L, Li E. Deep proteome profiling promotes whole proteome characterization and drug discovery for esophageal squamous cell carcinoma. Cancer Biol Med. 2022;19:273–7.

45. Shanabag A, Armand J, Son E, Yang HW. Targeting CDK4/6 in breast cancer. Exp Mol Med. 2025;57:312–22.

46. Wander SA, Cohen O, Gong X, Johnson GN, Buendia-Buendia JE, Lloyd MR, et al. The Genomic Landscape of Intrinsic and Acquired Resistance to Cyclin-Dependent Kinase 4/6 Inhibitors in Patients with Hormone Receptor–Positive Metastatic Breast Cancer. Cancer Discov. 2020;10:1174–93.

47. Soria-Bretones I, Thu KL, Silvester J, Cruickshank J, Ghamrasni SE, Ba-alawi W, et al. The spindle assembly checkpoint is a therapeutic vulnerability of CDK4/6 inhibitor–resistant ER+ breast cancer with mitotic aberrations. Sci Adv. 2022;8:eabq4293.

48. Ivanov A, Pawlikowski J, Manoharan I, van Tuyn J, Nelson DM, Rai TS, et al. Lysosome-mediated processing of chromatin in senescence. J Cell Biol. 2013;202:129–43.

49. Okada Y, Chikura S, Kimoto T, Iijima T. CDK4/6 inhibitor-induced bone marrow micronuclei might be caused by cell cycle arrest during erythropoiesis. Genes Environ. 2024;46:3.

50. Roggero CM, Ghosh AB, Devineni A, Ma S, Blatt E, Raj GV, et al. CDK4/6 inhibitors promote PARP1 degradation and synergize with PARP inhibitors in non-small cell lung cancer. Transl Oncol. 2025;52:102231.

51. MacDonald KM, Nicholson-Puthenveedu S, Tageldein MM, Khasnis S, Arrowsmith CH, Harding SM. Antecedent chromatin organization determines cGAS recruitment to ruptured micronuclei. Nat Commun. 2023;14:556.

52. Takaki T, Millar R, Hiley CT, Boulton SJ. Micronuclei induced by radiation, replication stress, or chromosome segregation errors do not activate cGAS-STING. Mol Cell. 2024;84:2203–2213.e5.

53. Zhang D, Liu Y, Zhu Y, Zhang Q, Guan H, Liu S, et al. A non-canonical cGAS–STING– PERK pathway facilitates the translational program critical for senescence and organ fibrosis. Nat Cell Biol. 2022;1–17.

54. Chen Q, Sun L, Chen ZJ. Regulation and function of the cGAS–STING pathway of cytosolic DNA sensing. Nat Immunol. 2016;17:1142–9.

55. Dvorkin S, Cambier S, Volkman HE, Stetson DB. New frontiers in the cGAS-STING intracellular DNA-sensing pathway. Immunity. 2024;57:718–30.

56. Singh V, Ubaid S, Kashif M, Singh T, Singh G, Pahwa R, et al. Role of inflammasomes in cancer immunity: mechanisms and therapeutic potential. J Exp Clin Cancer Res. 2025;44:109.

57. Almine JF, O’Hare CAJ, Dunphy G, Haga IR, Naik RJ, Atrih A, et al. IFI16 and cGAS cooperate in the activation of STING during DNA sensing in human keratinocytes. Nat Commun. 2017;8:14392.

58. Moresi F, Noto F, Vasudevan J, Bisteau X, Adriani G, Pavesi A. Microphysiological Systems in Cancer Research: Advancing Immunotherapy through Tumor Microenvironment-Integrated Organ-On-Chip Models. Adv Ther. 2025;

59. Vasudevan J, Vijayakumar R, Reales-Calderon JA, Lam MSY, Ow JR, Aw J, et al. In vitro integration of a functional vasculature to model endothelial regulation of chemotherapy and T-cell immunotherapy in liver cancer. Biomaterials. 2025;320:123175.

60. Goel S, DeCristo MJ, Watt AC, BrinJones H, Sceneay J, Li BB, et al. CDK4/6 inhibition triggers anti-tumour immunity. Nature. 2017;548:471–5.

61. Deng J, Wang ES, Jenkins RW, Li S, Dries R, Yates K, et al. CDK4/6 Inhibition Augments Anti-Tumor Immunity by Enhancing T Cell Activation. Cancer Discov. 2017;8:CD-17-0915.

62. Salvador-Barbero B, Álvarez-Fernández M, Zapatero-Solana E, Bakkali AE, Menéndez M del C, López-Casas PP, et al. CDK4/6 Inhibitors Impair Recovery from Cytotoxic Chemotherapy in Pancreatic Adenocarcinoma. Cancer Cell. 2020;37:340–353.e6.

63. Li K, Liu C, Sui X, Li C, Zhang T, Zhao T, et al. An organoid co-culture model for probing systemic anti-tumor immunity in lung cancer. Cell Stem Cell. 2025;

