## Supplementary figures and images for "CDK4/6 Inhibition Uncovers Subtype-Specific Vulnerabilities and Immune-Related Responses in Esophageal Squamous Cell Carcinoma"

### Supplemental Figure 1

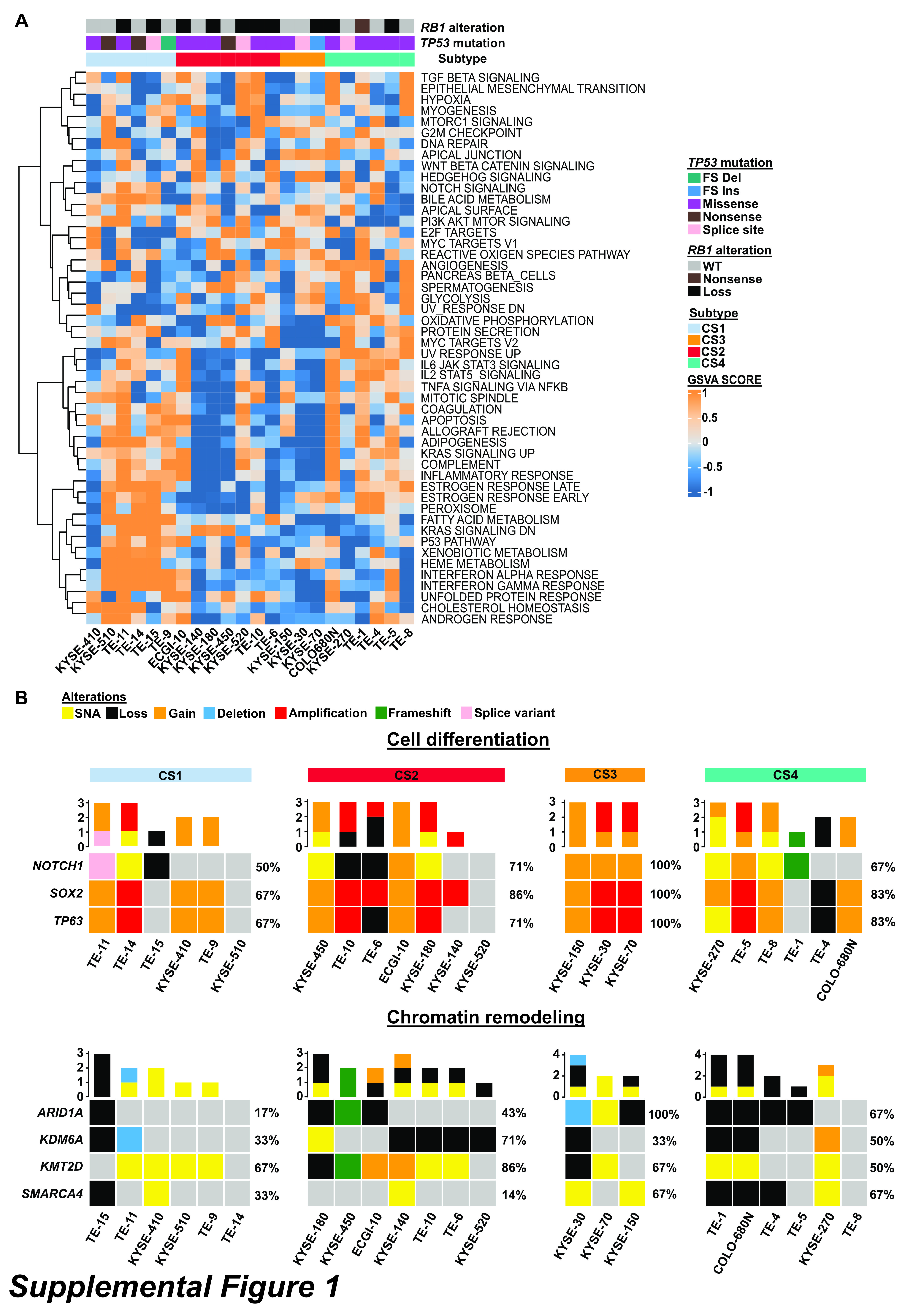

### Supplemental Figure 2

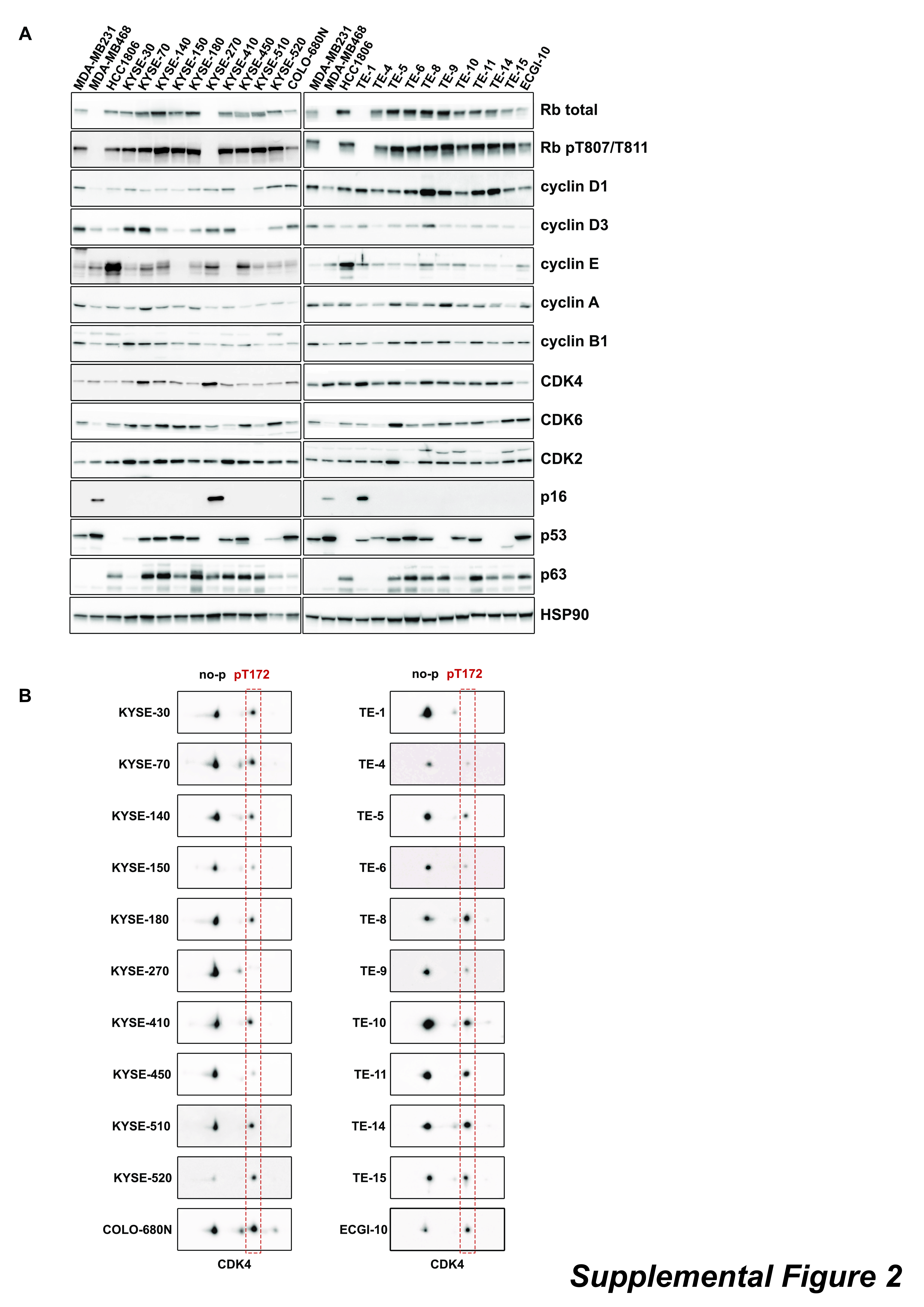

### Supplemental Figure 3

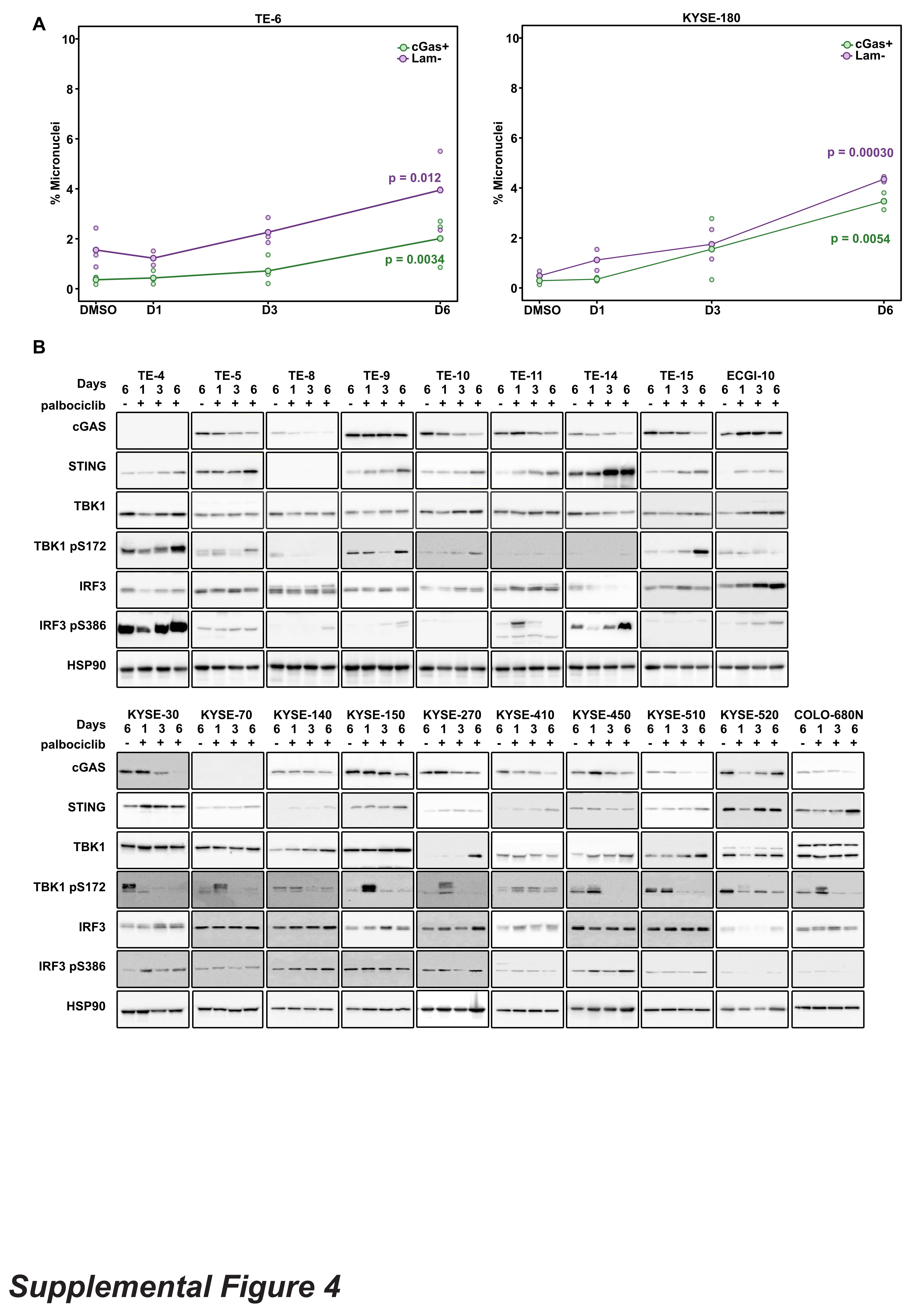

### Supplemental Figure 4

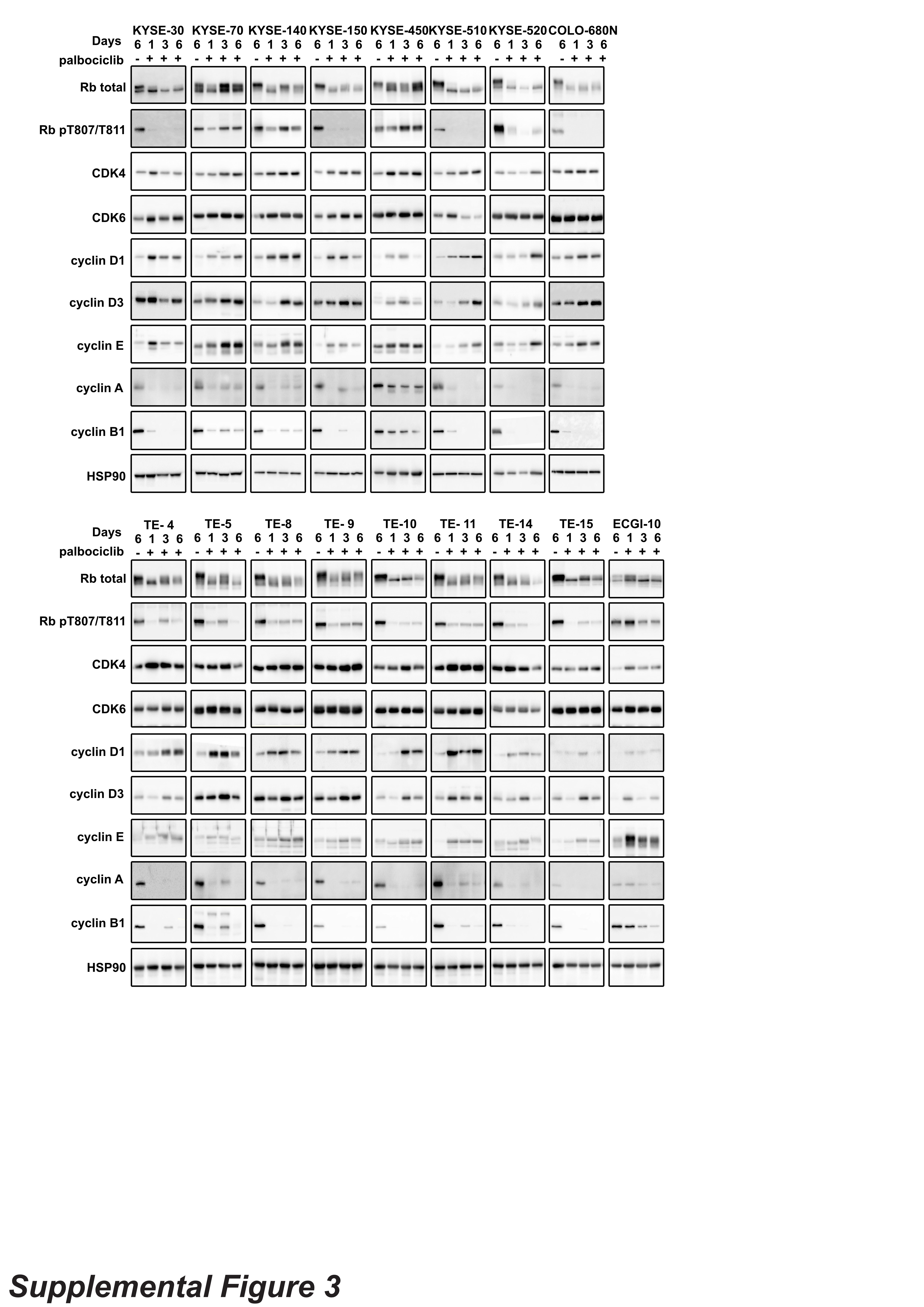
